# A transcontinental experiment elucidates (mal)adaptation of a cosmopolitan plant to climate in space and time

**DOI:** 10.1101/2024.09.16.613311

**Authors:** Lucas J. Albano, Cristina C. Bastias, Aurélien Estarague, Brandon T. Hendrickson, Simon G. Innes, Nevada King, Courtney M. Patterson, Amelia Tudoran, François Vasseur, Adriana Puentes, Cyrille Violle, Nicholas J. Kooyers, Marc T. J. Johnson

**Affiliations:** Department of Ecology and Evolutionary Biology, University of Toronto, Toronto, ON, Canada; Department of Biology, University of Toronto Mississauga, Mississauga, ON, Canada; Ecology Area, Department of Botany, Ecology and Plant Physiology, Campus de Rabanales, Córdoba University, Córdoba, Andalusia, Spain; CEFE, Université Montpellier, CNRS, EPHE, IRD, Montpellier, Occitanie, France; Department of Biology, University of Louisiana at Lafayette, Lafayette, LA, USA; Department of Ecology, Swedish University of Agricultural Sciences, Uppsala, Uppland, Sweden

**Author notes:** **Correspondence Author:** Lucas J. Albano, Current address: Department of Plant & Microbial Biology, North Carolina State University, Raleigh, NC, United States, 27695. These authors co-supervised the project (Adriana Puentes, Cyrille Violle, Nicholas J. Kooyers, Marc T. J. Johnson).

**Keywords:** Adaptation lag, climatic distance, common garden, cyanogenesis, environmental variation, introduced range, non-native species, local adaptation, native range, provenance study, *Trifolium repens*, white clover

## Abstract

Climate change and the global spread of non-native species are two of the most significant threats to biodiversity and ecosystem function. Both these phenomena subject populations to novel conditions, either in space (species introductions) or in time (climate change), yet the role of adaptation in how populations respond to these rapid environmental shifts is poorly understood. We conducted a large-scale trans-continental common garden experiment using white clover (*Trifolium repens*, Fabaceae) to test whether adaptive evolution to spatiotemporal variation in climate could contribute to the ecological success of one of the most widespread plant species in the world. Individuals from 96 populations of *Trifolium repens* (white clover) from both its native (Europe) and introduced (North America) ranges were planted into four experimental common gardens located in northern (Uppsala, Sweden) and southern (Montpellier, France) Europe, and northern (Mississauga, Canada) and southern (Lafayette, USA) North America. We recorded plant sexual and clonal fitness in each common garden and assessed whether the strength of local adaptation differed between the native and introduced ranges and whether populations are rapidly adapting to climate change. Results show that local adaptation was only evident when populations were transplanted into common gardens located in the same range (native or introduced) from which they originated and was driven by stronger selection (due to climatic factors rather than herbivory) at lower latitudes in both ranges. Our results indicate rapid local adaptation across a large latitudinal gradient in introduced *T. repens* populations, along with an associated adaptation cost when transplanted back into the native range. We also find evidence of an adaptation lag in the northern common garden in the introduced range, with plants from historically warmer climates exhibiting the greatest fitness. These findings support two major conclusions: 1) white clover can rapidly adapt to spatial variation in climate in its introduced range as well as the native range, and 2) despite rapid adaptation to novel environments, introduced white clover populations are not keeping pace with rapid climate change. Overall, our results provide insight into the role of adaptation in facilitating the ecological success of non-native species in a rapidly changing world.

**Open Research Statement:** Data are provided for peer review. All data involved in this study is available on the GitHub page for LJA (https://github.com/ljalbano/transcontinental_common_garden).

## Introduction

Natural selection results in populations evolving traits that are advantageous to the environmental conditions of a particular location, a phenomenon known as local adaptation (Turesson, 1922; Kawecki and Ebert, 2004; Hereford, 2009). A fundamental aspect of local adaptation is that variation in biotic and abiotic factors leads to changes in the relative fitness of genotypes between environments (Gillespie and Turelli, 1989; Kawecki and Ebert, 2004; Lee and Mitchell-Olds, 2011). Such genotype-by-environment interactions can be elucidated by reciprocal transplant experiments, which are challenging to conduct, but vital in demonstrating fitness trade-offs, whereby genotypes exhibit higher relative fitness in their local environment and lower fitness in other environments (Gillespie and Turelli, 1989; Hereford, 2009; Leimu and Fischer, 2008; Savolainen et al., 2013). Human activities are exposing populations to rapidly changing novel environments, both across space through species introductions (Pyšek et al., 2020), and over time through anthropogenically mediated climate change (Hoffmann and Sgrò, 2011). Understanding how populations simultaneously adapt to novel biotic and abiotic environments over space and time is an important problem that is poorly resolved (Hendry, 2017; Siepielski et al., 2017). Addressing this problem is necessary to determine how increasingly pervasive anthropogenic disturbances alter the ecology and evolution of native and introduced species (Parmesan, 2006; Fugère and Hendry, 2018).

Introduced populations are frequently subjected to shifts in selection pressure compared to the native environment to which they are locally adapted (Erfmeier, 2013; Colautti and Lau, 2015; Lau and terHorst, 2015). This shift in selection pressure in either the biotic environment (e.g., novel species interactions) or the abiotic environment (e.g., climatic factors) could inhibit the establishment and spread of an introduced population (Erfmeier, 2013; Colautti and Lau, 2015; Lau and terHorst, 2015). Additionally, inherent characteristics of introduced populations, such as small population size and low genetic diversity, could limit their ability to respond to novel selection pressures, with Allee effects and genetic bottlenecks potentially resulting in mate limitation and inbreeding depression (Tsutsui et al., 2000; Kolar and Lodge, 2001; Allendorf and Lundquist, 2003; Taylor and Hastings, 2005; Burns et al., 2011; Lau and terHorst, 2015; Pannell et al., 2015; Estoup et al., 2016). In most cases, the ecological and/or genetic constraints experienced by introduced populations are sufficient to result in the failure of those populations to establish and their eventual extirpation (Kolar and Lodge, 2001; Jeschke and Pyšek, 2018).

Despite the challenges presented by introduction to novel environments across space, introduced populations still often become invasive, experiencing substantial ecological opportunity if constraints can be overcome. For example, ecological constraints can be overcome if novel selection regimes in introduced environments are either benign or even favourable to a non-native population (Erfmeier, 2013; Colautti and Lau, 2015). This is particularly true when an introduced population has an advantage over its new natural enemies or competitors compared to in its native range (Blossey and Nötzold, 1995; Callaway and Ridenour, 2004; Müller-Schärer et al., 2004; Callaway et al., 2022). It is also possible that introduced populations do not exhibit strong evidence of genetic constraints to begin with (i.e., the genetic paradox of invasions), because reductions in genetic variation were not substantial enough to limit the ability of a population to respond to novel selection pressures to begin with (Allendorf and Lundquist, 2003; Maron et al., 2004; Dlugosch and Parker, 2008a; Dlugosch and Parker, 2008b; Colautti and Barrett, 2013; Estoup et al., 2016; van Boheemen et al., 2017; Battlay et al., 2024). In fact, when introduced populations experience strong selection and sufficient genetic variation in novel environments, it could facilitate rapid adaptation to those novel environments, which could in turn, contribute to the success of a non-native species (Maron et al., 2004; Dlugosch and Parker, 2008a; Colautti and Barrett, 2013; Colautti and Lau, 2015; Simón-Porcar et al., 2021; but see Genton et al., 2005). However, the extent to which rapid local adaptation to novel biotic and abiotic conditions can facilitate the success of a widespread non-native species across large environmental gradients remains poorly understood.

While there are examples in which introduced populations have rapidly adapted to novel environments across space, it is important to consider that these novel environments may also be changing through time. Temporal shifts in a local environment can cause selection pressures to fluctuate, which could result in traits that were previously adapted to that environment becoming maladaptive (Wilczek et al., 2014; Aguirre-Liguori et al., 2019; Exposito-Alonso et al., 2019; Anderson and Wadgymar, 2020; Bontrager et al., 2020; Capblancq et al., 2020). If the rate of adaptation is exceeded by the rate of environmental change, this disruption of local adaptation is likely to result in reduced fitness (i.e., maladaptation) of populations (St Clair and Howe, 2007; Atkins and Travis, 2010; Wilczek et al., 2014; Aguirre-Liguori et al., 2019; Capblancq et al., 2020). Recent global warming may result in populations becoming maladapted to their local climate conditions, whereas populations from lower latitudes or elevations (i.e., warmer origins) become better suited to environmental conditions at higher latitudes or elevations; a phenomenon known as an adaptation lag (Wilczek et al., 2014; Kooyers et al., 2019; Anderson and Wadgymar, 2020). Currently, the frequency and magnitude of adaptation lags are poorly understood (Anderson and Song, 2020).

If non-native species rapidly adapt to novel environments across space, similar rapid adaptation to climate change through time could facilitate their continued range expansion, persistence, and pervasiveness (Byers, 2002; Maron et al., 2004; Colautti and Barrett, 2013; Oduor et al., 2016). In fact, non-native species are often better at climate tracking and more resistant to climate warming than native species, which could contribute to their success as invaders (Wolkovich et al., 2013). However, the interplay between local adaptation, species introductions, and anthropogenically mediated climate change remains an understudied, yet critically important, aspect of understanding the ongoing human-induced biodiversity crisis (IPBES, 2019). This problem is best addressed through large field experiments testing local adaptation in both the native and introduced ranges of widespread non-native species, alongside elucidating the environmental conditions that may result in rapid adaptation and success in a novel environment (Colautti and Lau, 2015; Anderson and Song, 2020).

Plant-herbivore interactions have been highlighted as key selective factors that determine patterns of local adaptation, species responses to environmental change, and biological invasions. Spatial variation in both biotic and abiotic factors are likely to lead to spatial variation in plant defense traits by affecting the costs and benefits involved in growth-defense trade-offs (Anstett et al., 2018; Baskett and Schemske, 2018; Moreira et al., 2018). Local herbivore communities impose selection through preferential feeding on plants with lower resistance, influencing the benefits of plant defenses in areas of particularly high herbivore pressure(Ehrlich and Raven, 1964; Coley and Barone, 1996; Agrawal, 2007; Johnson and Rasmann, 2011; Züst et al., 2012). In turn, the metabolic costs of defenses can be substantial, while abiotic environments and the stressors they impose can influence the availability of resources to plants, exacerbating defense costs (Coley et al., 1985; Herms and Mattson, 1996; Fine et al., 2006; Hahn et al., 2019). Furthermore, it is increasingly recognized that anthropogenic activity can influence the evolution of plant defenses (Bidart-Bouzat and Imeh-Nathaniel, 2008; Massad and Dyer, 2010; Miles et al., 2019). Climate warming could result in increased herbivore feeding and/or increased plant defenses (Dostálek et al., 2020; Lemoine et al., 2013) or could contribute to success of a non-native species by decoupling locally adapted plant-herbivore interactions (Adams et al., 2009; Costan et al., 2022; but see Agrawal and Kotanen, 2003). It is therefore important to understand how populations are adapting to their local biotic and abiotic environments and how anthropogenic factors can affect adaptation.

An ideal study system to test questions surrounding adaptation to novel environments post-introduction and in response to climate change would be a widespread species with known agents of biotic and abiotic selection across space and over time that can drive adaptation in the native and introduced ranges. *Trifolium repens* is an allopolyploid perennial legume that is native to Eurasia and now has a cosmopolitan distribution, following introduction to inhabited continents throughout the world, including North America (Daday, 1958; Kjærgaard, 2003; Griffiths et al., 2019). Mechanisms of biotic and abiotic selection in *T. repens* have been studied for over 70 years. In *T. repens*, clines in a prominent chemical defense mechanism, hydrogen cyanide (HCN), have been identified across many types of environmental gradients (Daday, 1954a; Daday, 1954b, Daday, 1958; Till, 1987; Kooyers and Olsen, 2013; Kooyers et al., 2014; Thompson et al., 2016), providing opportunities to test hypotheses surrounding the involvement of biotic factors in adaptation to environmental change.

To understand adaptation to environmental variation in a non-native species over space and time, we conducted a trans-continental common garden field experiment across latitudinal gradients in both the native (Europe) and introduced (North America) ranges of *T. repens*. We first address four research questions focused on adaptation in *T. repens* to biotic and abiotic environmental variation in space. (Q1) Are *T. repens* populations locally adapted to their climate of origin and does the strength of local adaptation differ across space? This question is a prerequisite condition before any comparisons can be made regarding local adaptation between the native and introduced ranges. (Q2) Does *T. repens* exhibit stronger local adaptation in its native range than in its introduced range?. (Q3) What are the roles of plant defense and herbivory (i.e., biotic factors) in driving local adaptation across space? (Q4) What bioclimatic variables (i.e., abiotic factors) have the strongest impact on fitness across space (latitudes and ranges) in *T. repens*? We then focus on whether *T. repens* can adapt to climate change through time using a space-for-time approach to analyzing a subset of the common garden experimental data (Pickett, 1989; Fournier-Level et al., 2011), asking: (Q5) Are there adaptation lags to climate change in *T. repens*, such that populations originating from warmer and/or drier climates exhibit greater fitness than local populations? This study is among the first to investigate the ability of an introduced species to rapidly adapt to novel environments in space and time simultaneously. Our results elucidate the potential for rapid adaptation to contribute to initial and continued ecological success of non-native species in the face of ongoing environmental change.

## Methods

### Study system

*Trifolium repens* has been a model for the study of plant defense based on its polymorphism for the production of the potent chemical defense hydrogen cyanide (HCN; Hughes, 1991). In fact, the ability to produce HCN as an antiherbivore defense may have contributed to the range expansion of *T. repens*, allowing it to thrive in human-disturbed habitats such as lawns, fields, and roadsides in many regions throughout the world (Burdon, 1983; Kjærgaard, 2003; Olsen et al., 2021). Cyanogenesis occurs through the hydrolyzation of cyanogenic glucosides by the enzyme linamarase. In *T. repens*, the presence or absence of cyanogenic glucosides, linamarin and lotaustralin, is controlled by three linked genes in their metabolic pathway (*CYP79D15*, *CYP736A187*, and *UGT85K17*; referred to collectively as *Ac*/*ac*), while the presence or absence of the enzyme linamarase is controlled by the gene *Li*/*li* (Olsen et al., 2007; Olsen and Small, 2018). Thus, four phenotypes exist for HCN and the two metabolic components required for its production: one in which HCN is constitutively produced (AcLi) and three that are acyanogenic and therefore lack the defense entirely (acLi, Acli, acli; Corkill, 1940; Atwood and Sullivan, 1943).

Cyanogenesis is widely considered an adaptive polymorphism because *T. repens* exhibits clines in the presence of HCN across numerous types of environmental gradients (Daday, 1954a; Daday, 1954b; Daday 1958; Brighton and Horne, 1977; Thompson et al., 2016; Innes et al., 2022; Santangelo et al., 2022). In general, dominant *Ac* and *Li* alleles (and thus the ability to produce HCN) are fixed in warmer climates, while recessive *ac* and *li* alleles are fixed in colder climates (Daday, 1954a; Daday, 1954b; Daday, 1965). Latitudinal cyanogenesis clines have been identified in the native European range of *T. repens* (Daday, 1954a), as well as its introduced ranges, such as North America (Daday, 1965). However, clines within introduced ranges as generally shallower, possibly indicating weaker local adaptation post-introduction, which occurred less than 400 years ago in the case of North America (Daday, 1958; Kjærgaard, 2003; Kooyers and Olsen, 2012; Innes et al., 2022). One hypothesis to explain cyanogenesis clines is a gradient in herbivory, with greater diversity and abundance of herbivores in warmer climates resulting in stronger selection favouring cyanogenic plants (Angseesing, 1974; Dirzo and Harper, 1982; Kakes, 1989). Alternatively, empirical studies across various environmental gradients have identified temperature, soil moisture, and soil nutrients as abiotic factors that may impose selection on cyanogenesis or its metabolic components to influence cline formation (Daday, 1954a; Daday, 1965; Gleadow and Woodrow, 2002; Kooyers et al., 2014; Kooyers et al., 2018). Overall, despite 70 years of research on the drivers of cyanogenesis clines in *T. repens*, there remains little consensus on how variation in environmental conditions influence selection on cyanogenesis in *T. repens*, or if cyanogenesis is primarily responsible for rapid local adaptation to novel environments (Daday, 1954a; Daday, 1965; Hughes, 1991; Kooyers and Olsen, 2013; Kooyers et al., 2014; Wright et al., 2018; Wright et al., 2021; Albano and Johnson, 2023; Kuo et al., 2024). Our study aims to resolve this uncertainty by directly testing the strength of local adaptation and its biotic and abiotic drivers in both the native and introduced ranges of *T. repens*. *Sampled populations and common garden sites*

We addressed our research questions using four common gardens distributed across northern (Uppsala, Sweden) and southern (Montpellier, France) regions of the native European range of *T. repens*, and northern (Mississauga, ON, Canada) and southern (Lafayette, LA, USA) regions of the North American introduced range (**Figure 1**). Seeds were collected from 49 natural *T. repens* populations scattered across the native European range spanning ∼27° of latitude, and 47 natural *T. repens* populations along a 2500 km north-south transect spanning ∼21° of latitude in the introduced North American range (**Figure 1**). Temperature and latitude are highly correlated in eastern North America but less directly associated in Europe, with temperatures varying across both north-south and east-west gradients, which led to the more diffuse sampling approach in Europe and the transect sampling approach in North America. To minimize maternal and epigenetic effects, we hand-pollinated plants germinated from field-collected seeds in a growth chamber environment (25°C day, 15°C night, 14 h:10 h light:dark cycle; Conviron MTPS, Winnipeg, MB, Canada) to create an outbred F1 cohort of seeds for each population. F1 generation seeds were germinated and established for four weeks in peat pots (8 cm diameter, 591 mL volume) in uniform growth chamber or greenhouse conditions near each common garden site. Five replicate individuals from each population were transplanted in peat pots directly into the natural soil of a typical cultivated lawn at each common garden site. Inconsistent germination success resulted in some populations not being included in some common garden locations, and in very limited cases, populations had four or six individuals planted. The resulting total number of plants per garden was 465, 465, 458, and 470 in Mississauga, Lafayette, Uppsala, and Montpellier, respectively. Plants were arranged in rows and columns with 1 m spacing, and black landscaping fabric was used to cover the entirety of the lawn to prevent the rooting of stolons into the natural soil between plants. Holes (12 cm diameter) were cut in the fabric for each individual plant, leaving a 2 cm ring of natural vegetation around each plant to allow for natural competition. This vegetation was trimmed monthly throughout the experiment. Planting dates matched growing seasons in each location. The Lafayette and Mississauga common gardens were established in April and May 2020, respectively, while the Montpellier and Uppsala common gardens were postponed due to COVID-19 restrictions in Europe, leading to their establishment in March and May 2021, respectively. Plants were given supplemental water for the first two weeks of the experiment and intermittently after the first two weeks when necessary to avoid excessive mortality. Plants that died within the first two weeks of the experiment were deemed to have died due to transplantation stress and were replaced using extra germinated individuals from the same population (Mississauga: 3; Lafayette: 32; Uppsala: 72; Montpellier: 84). Plants that died after two weeks were deemed to have died due to the local environmental conditions at each common garden site and were not replaced. Each common garden was maintained for two full growing seasons.

**Figure 1.**
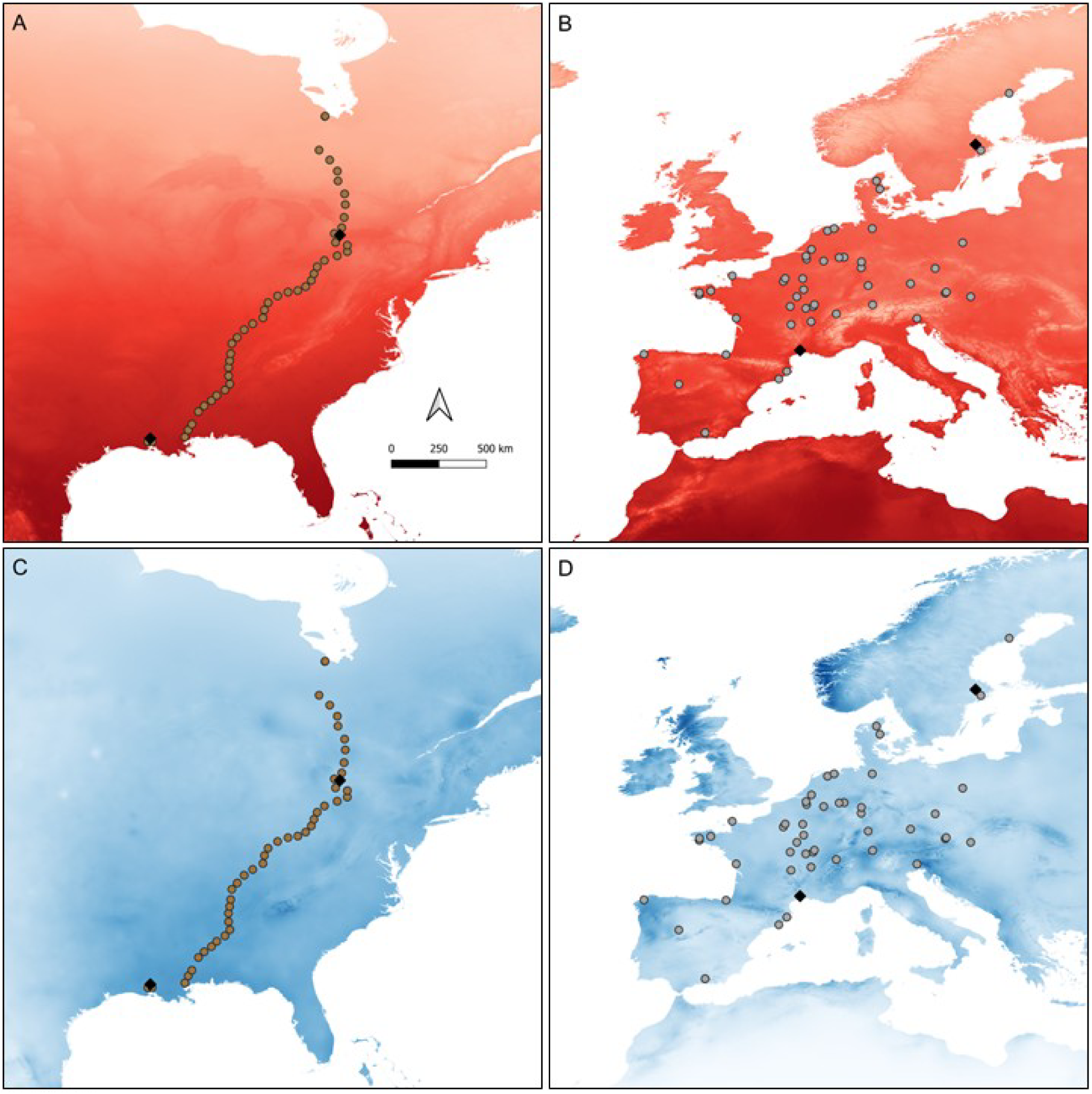
Maps showing populations of *T. repens* sampled from the (**A**, **C**) introduced range (North America, circles) and (**B**, **D**) native range (Europe, grey circles), as well as the four common garden locations (black diamonds). Map panels **A** and **B** are set on a backdrop of colour-graded historical (1970-2000) mean annual temperature (BIO1; with darker red shades representing warmer climates). Map panels **C** and **D** are set on a backdrop of colour-graded historical (1970-2000) annual precipitation (BIO12; with darker blue shades representing wetter climates). Coordinates of common garden locations are 42°32’40” N, 79°39’38” W (Mississauga, Ontario, Canada), 30°18’23” N, 92°00’33” W (Lafayette, Louisiana, USA), 59°49’08” N, 17°38’49” W (Uppsala, Sweden), and 43°38’16” N, 3°51’43” W (Montpellier, France). All climate data was extracted from WorldClim (Version 2.1, February 2023).

### Tissue collection, herbivory estimates, and measures of fitness

After 1-2 months of growth, one to three fully-expanded leaves were harvested from each surviving individual and frozen at −20°C. These leaves were later freeze-dried and used for DNA extraction, followed by low coverage whole genome sequencing to predict the presence of *Ac/ac* and *Li/li* alleles (see the **Supplementary Methods** section of the **Supporting Information**). In total, 586 individuals (Mississauga: 302, Lafayette: 99, Uppsala: 78, Montpellier: 107) were successfully sequenced and characterized as possessing either dominant or recessive alleles at *Ac/ac* and *Li/li.* The remaining 1272 individuals were not genotyped because they either died before tissue could be harvested or genomic library preparations were unsuccessful. We exposed plants to the natural environment of each common garden at the earliest life stage possible to capture potentially important selection on seedlings (Agrawal et al., 2012). As a result, the experiment experienced high mortality of young individuals, which is common and a natural part of the biology of *T. repens* and other plants (Escudero et al., 2000; Moles and Westoby, 2004; Agrawal et al., 2012). This mortality prevented many individuals from being included in tissue collection. Visual herbivory estimates (% leaf area consumed) were conducted monthly following the methods of Johnson et al. (2016). Herbivory was analyzed for the month exhibiting peak herbivory (highest average % leaf area consumed) within each garden. Multiple components of plant fitness were measured throughout the growing seasons at each location. Survival and onset of flowering were assessed weekly during the first growing season. Mature fruits were counted and collected weekly, and seeds were separated using a thresher (Precision Machine Co., Lincoln, NE, USA) to obtain total seed set mass for each plant across both growing seasons of the experiment. Photos were taken of each plant monthly during the first growing season at each common garden location and assessed for plant area using Easy Leaf Area (Version 2.0; Easlon and Bloom, 2014). The maximum plant area was then determined and the growth rate across the first growing season was calculated as (ln[A_max_] – ln[A_1_])/(t_max_ – t_1_), where A_max_ represents the maximum plant area, A_1_ represents the initial plant area (i.e., after one month of growth in the common garden), and (t_max_ – t_1_) represents the number of days between photos taken for maximum plant area and initial plant area.

### Calculating climatic distance

Six bioclimatic variables were extracted from WorldClim (Version 2.1, February 2023) at a resolution of 30 seconds for each original *T. repens* population and common garden site (O’Donnell and Ignizio, 2012). The variables included were mean annual temperature (“BIO1”; MAT), maximum temperature of the warmest month (“BIO5”), minimum temperature of the coldest month (“BIO6”), annual precipitation (“BIO12”), precipitation of the wettest month (“BIO13”), and precipitation of the driest month (“BIO14”). The extracted values of each bioclimatic variable for each common garden location can be found in **Table S1**. Values for all six variables were z-score standardized to allow for analysis on an equivalent scale. A principal component analysis was performed using the *princomp* function in R, with a scree plot to guide the decision to retain principal components 1, 2, and 3, which explained 65.8%, 22.1%, and 9.1% of the variation, respectively (cumulatively explaining 97% of variation; **Figure S1**).

Coordinates for each location of origin and each common garden location from all three dimensions were then extracted using the *get_pca_ind* function in the *factoextra* package (version 1.0.7; Kassambara and Mundt, 2020), and then plotted on a 3D graph (**Figure S2**). The weighted Euclidean distance between points in 3D space was calculated using the formula:

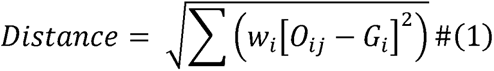

Where *w_i_* represents the proportion of variance explained by the *i*th dimension, *O_ij_* represents the value of the *j*th population of origin in the *i*th dimension, and *G_i_* represents the value of a given common garden in the *i*th dimension. We used this weighted Euclidean distance as a measure of the climatic distance between the location of each population of origin and each common garden location based on our six bioclimatic variables of interest (Wright et al., 2018; Samis et al., 2019). This measure of climatic distance was then used as a predictor variable of plant fitness in our statistical analyses. Bivariate correlations among individual climatic variables and with climatic distance in each common garden are reported in **Table S2**. We also tested average climatic distances among sampled populations within each continent transplanted into common gardens on each continent using the following model (R function *lm*):

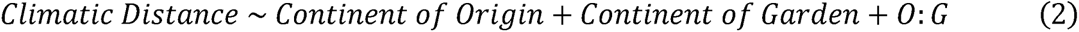

Climatic distance varied based on continent of origin (*F*_1,1854_ = 93.849, *P* < 0.001), continent of garden (*F*_1,1854_ = 65.960, *P* < 0.001), and their interaction (*F*_1,1854_ = 142.491, *P* < 0.001), with European populations transplanted into European common gardens generally experiencing the smallest climatic distances between environment of origin and the common garden environment. *Statistical analysis*

All statistical procedures and analyses were conducted in R Version 4.1.2 (2021) and RStudio Version 2022.12.0 (R Core Development Team, 2022). Seven response variables were assessed as components of fitness: i) plant survival in the first growing season, ii) presence/absence of flowers during either growing season, iii) presence/absence of seeds during either growing season, iv) number of flower heads produced across both growing seasons, v) seed set mass produced across both growing seasons, vi) maximum plant area in the first growing season, and vii) growth rate in the first growing season. Herbivory during the first growing season was also measured as an additional non-fitness response variable. The number of flower heads and seed set mass were both zero-inflated, which led to the use of a two-step modelling approach as described previously (Albano and Johnson, 2023). Briefly, both variables were first analyzed in binomial models to assess whether plants produced flowers or seeds. Non-zero flower head numbers and seed set masses were then subsequently analyzed as separate fitness components. All linear mixed effects models were conducted using the *glmmTMB* function in the *glmmTMB* package (version 1.1.5; Brooks et al., 2017). All test statistics and *P* values were calculated using the *Anova* function in the *car* package (version 3.0-12; Fox and Weisberg, 2019) with Type II sums-of-squares. We fit three binary fitness measures (i.e., plant survival, presence/absence of flowers, presence/absence of seeds) to a binomial distribution. All quantitative measures of fitness (i.e., number of flower heads, seed set mass, maximum plant area, growth rate) and herbivory were modeled using a normal distribution, with all transformations necessary to meet assumptions of normality and homogeneity of variance found in **Table S3** and sample sizes for each response variable and each model found in **Table S4**. Slope estimates and their standard errors were calculated using the *emtrends* function in the *emmeans* package (version 1.10.4; Lenth, 2024).

### Variation in the strength of local adaptation between common garden locations

To determine whether *T. repens* populations exhibit local adaptation and to assess the strength of local adaptation at each garden location (Q1), all response variables were analyzed using the following structure:

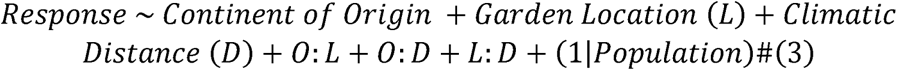

Continent of origin (North America or Europe), garden location (Mississauga, Lafayette, Uppsala, or Montpellier), and climatic distance (the combined measure of six bioclimatic variables of interest described above) were included as fixed effects, along with all two- and three-way interactions. Population was included as a random effect to account for non-independence of replicates within each population. The strength of local adaptation refers to the magnitude of the fitness advantage for local populations compared to populations transplanted at an increasingly greater climatic distance from their origin. We therefore interpret the slope of the relationship between climatic distance and fitness as being representative of the strength of local adaptation within a given common garden (a negative slope of greater magnitude represents greater strength of local adaptation). An interaction between continent of origin and climatic distance would indicate that the strength of local adaptation varies based on whether populations originated in the native or introduced range. An interaction between garden location and climatic distance would indicate that the strength of local adaptation varies based on the common environment in which it is being tested.

### Variation in the strength of local adaptation between ranges

To assess the strength of local adaptation between the native and introduced ranges of *T. repens* (Q2), the analyses of the previous subsection were repeated, with the continent of the common garden (North America or Europe) replacing the location of the common garden in the models:

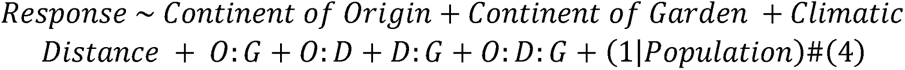

Interpretation of this model is similar to the model above, with an interaction between continent of origin and climatic distance indicating that the strength of local adaptation varies based on whether populations originated in the native or introduced range. An interaction between continent of garden and climatic distance would indicate that the strength of local adaptation varies based on the continent of the common environment in which it is being tested. In this model, we are particularly interested in a three-way interaction between continent of origin, continent of garden, and climatic distance because it would indicate that the strength of local adaptation varies based not only on the range from which populations originated but also the range in which they are being tested. In other words, this model aims to test whether locally adapted European populations maintain signatures of local adaptation in North America and vice versa.

### The roles of cyanogenesis and herbivory in local adaptation

The presence of cyanogenesis clines across latitude on both continents was determined separately for *Ac/ac* and *Li/li* by assessing the proportion of plants in each population containing at least one dominant *Ac* or *Li* allele using the following model structure fit to a binomial distribution:

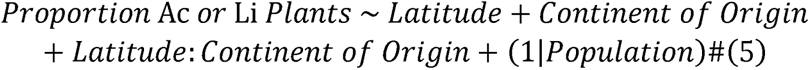

An interaction between latitude and continent of origin would indicate a difference in the slope of clines in the presence of *Ac* and/or *Li* based on the range in which those clines are occurring (native European or introduced North American).

The effect of cyanotype (AcLi, Acli, acLi, or acli) on local adaptation (Q3) was assessed for all eight response variables using the following model structure:

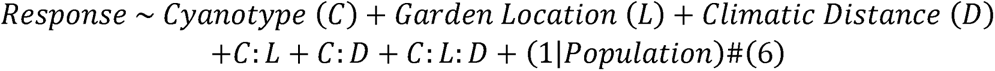

An interaction between cyanotype and climatic distance would indicate that cyanogenesis or its metabolic components contribute to the strength of local adaptation in *T. repens*. A three-way interaction between cyanotype, garden location, and climatic distance would indicate that selection for or against a given cyanotype in a particular common environment impacts the strength of local adaptation in that environment. This might be expected due to consistently observed latitudinal cyanogenesis clines across multiple types of environmental gradients in *T. repens*, which indicate selective advantages for particular cyanotypes depending on the location in which they are found, and the biotic and abiotic factors imposing selection in that location.

### Bioclimatic variables as individual drivers of local adaptation

We also measured associations between the seven response variables and each of the six bioclimatic variables of interest individually by determining the difference in z-score standardized values for each variable between the location of each population of origin and each common garden location using the following model structure:

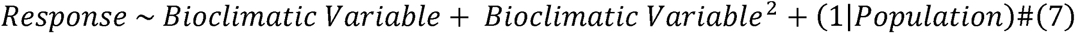

For this set of analyses, all common gardens were analyzed separately, and all quantitative (non-binomial) response variables were z-score standardized to allow for simultaneous direct comparisons of the relative strength of linear and quadratic predictor variables. Within each garden, we were therefore able to directly compare *P* values across all bioclimatic variables for both linear and quadratic fits (**Tables S5-S8**). For each response variable, we selected the most influential bioclimatic variable (i.e., the lowest *P*, but only when at least one *P* value was significant at a level of *P* < 0.1, highlighted in red in **Tables S5-S8**). Across all response variables in each garden, we then calculated the percentage of these most influential bioclimatic variables that were either temperature-based (BIO1, BIO5, or BIO6) or precipitation-based (BIO12, BIO13, and BIO14) and compared these percentages across the entire experiment, across latitudes, and across ranges (Q4). This approach was used to identify the most important abiotic variables driving local adaptation in general, as opposed to focusing on the results of any one particular test.

### Adaptation lags in response to climate warming

We sought to investigate the presence of adaptation lags using a space-for-time substitution (Q5). We performed an additional analysis using a similar model structure as before, but only using changes in raw values for MAT (i.e., not z-score standardized, hereafter referred to as ΔMAT) as the bioclimatic variable of interest.

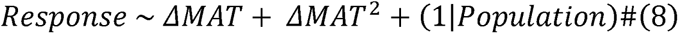

If the peak of the quadratic relationship in each model is approximately equal to zero, it would be considered evidence of local adaptation to MAT, because maximum fitness occurs in populations with the same mean annual temperature as the common garden site. If the peak of the quadratic relationship is instead shifted towards populations of warmer origins than are occurring at the common garden site (i.e., these warmer-adapted populations are outperforming the local populations), this would be considered evidence of an adaptation lag. We tested whether the shift of peaks away from a ΔMAT of zero and towards populations of warmer origins were significant by also fitting a similar model to above, but excluding the linear predictor:

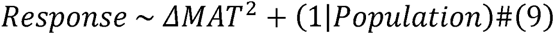

This additional model constrains the peak of the distribution to be centred around a ΔMAT of zero (approximating the outcome of local adaptation as opposed to an adaptation lag). We then compared the models with and without the linear predictor using the *anova* and *AIC* functions in R, with *P* values < 0.05 (along with ΔAIC values < 0) indicating a significant shift of the peak.

We focused our interpretation of this analysis based solely on the North American populations in the Mississauga common garden because the sampling design of North American populations along a transect and the location of the Mississauga garden within that sampled transect provide the most appropriate subset of the data to detect spatial lags. We include the results for European populations within the Uppsala common garden in **Table S9** for completeness, but the Uppsala common garden did not exhibit any trend in fitness based on climatic distance, precluding the possibility of an adaptation lag being detected. The Lafayette and Montpellier common gardens were excluded from this analysis because they are near the southern extremes of our sampled populations, precluding the ability to detect adaptation lags.

## Results

### Q1: Are T. repens populations locally adapted to their climate of origin and does the strength of local adaptation differ across space?

We found strong evidence of local adaptation, and the strength of local adaptation in *T. repens* varied among gardens for multiple fitness components. Climatic distance between each population of origin and the common garden locations strongly predicted all sexual fitness components and survival (**Table 1**). In general, populations tended to have higher fitness with lower climatic distance, indicating that populations of more similar climatic origins to the climate of each common garden were better adapted. The only response variables not showing evidence of local adaptation were the measures of vegetative growth used as proxies for clonal fitness (i.e., maximum plant area and growth rate). The interaction of common garden location and climatic distance affected the number of flower heads and seed set mass, but not other fitness components or herbivory (**Table 1**; **Figure S3**). In general, the strength of local adaptation, measured as the slope of the relationship between sexual fitness and climatic distance, was greater in the more southern common garden location on each continent (**Figure 2**). For the number of flower heads, the slope for Lafayette was 3.4× steeper than the slope for Mississauga and the slope for Montpellier was 349× steeper than the slope for Uppsala. Similarly, for seed set mass, the slope for Lafayette was 2.3× steeper than the slope for Mississauga and the slope for Montpellier was 384× steeper than the slope for Uppsala. The main effect of garden location affected all fitness components and herbivory, which reflects the large differences in environment between the gardens. By contrast, the main effect of continent of origin and its interactions with garden location and/or climatic distance did not affect any fitness component or herbivory (**Table 1**).

**Figure 2.**
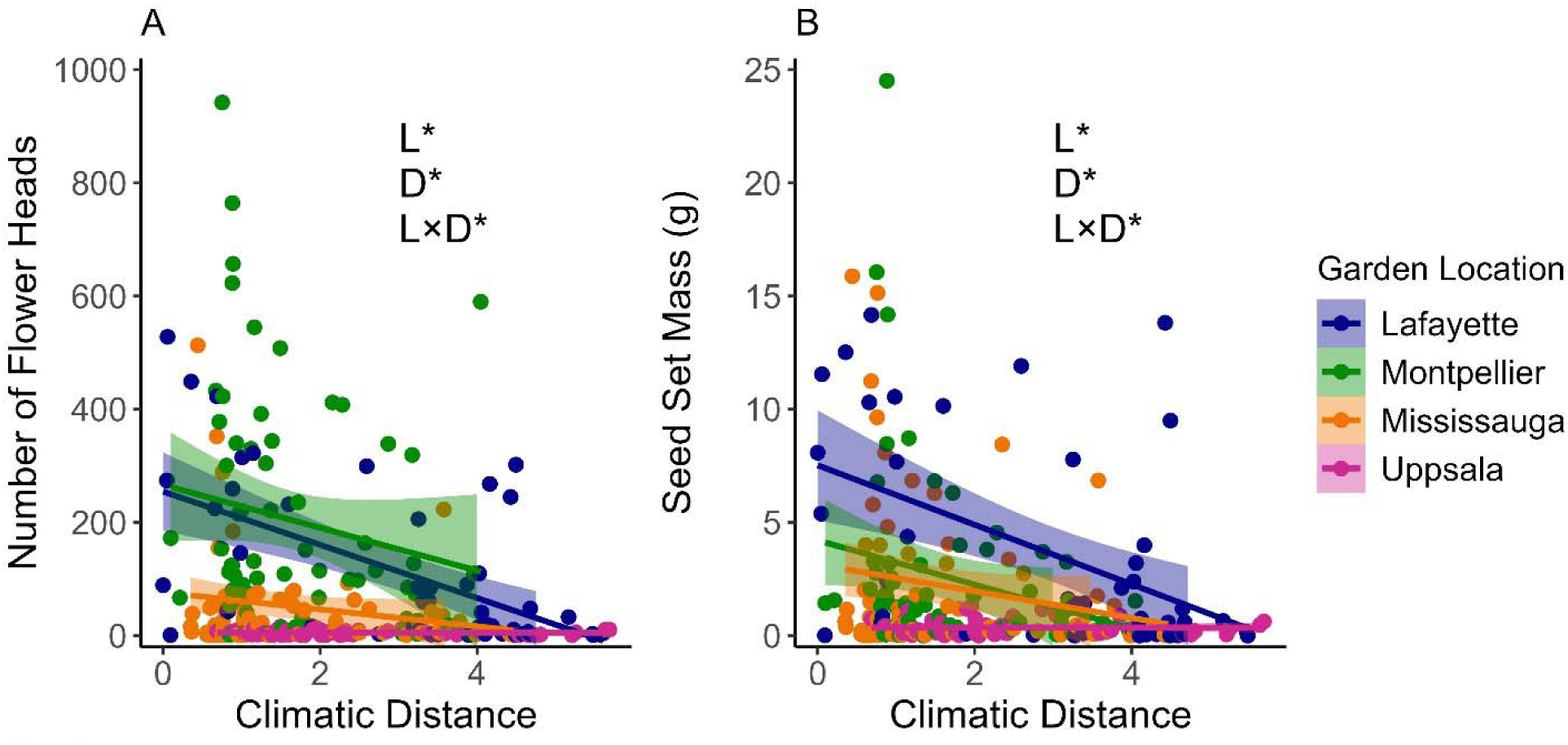
The effect of climatic distance and common garden location on fitness of *T. repens*. (**A**) Mean number of flower heads and (**B**) mean seed set mass of *T. repens* populations planted in two common gardens in North America (Lafayette in navy and Mississauga in orange) and two common gardens in Europe (Montpellier in green and Uppsala in magenta). Shaded areas represent 95% confidence intervals around each line, while asterisks represent *P* values of relevant fixed effects of Garden Location (L), Climatic Distance (D) and their interaction (\**P* < 0.05, \*\**P* < 0.01, \*\*\**P* < 0.001).

**Table 1.** Results of mixed effects models assessing how continent of origin, common garden location, and climatic distance affected multiple fitness components of *T. repens*.

| Predictor | <u>Plants Producing Flowers</u> |  |  | <u>Plants Producing Seeds</u> |  |  |
| --- | --- | --- | --- | --- | --- | --- |
| | $\chi^2$ | df | P | $\chi^2$ | df | P |
| Continent of Origin (O) | 0.377 | 1 | 0.539 | 0.271 | 1 | 0.603 |
| Garden Location (L) | **32.302** | **3** | **< 0.001** | **8.023** | **3** | **0.046** |
| Climatic Distance (D) | **6.308** | **1** | **0.012** | **7.197** | **1** | **0.007** |
| O × L | 0.773 | 3 | 0.856 | 0.724 | 3 | 0.868 |
| O × D | 0.299 | 1 | 0.584 | 0.876 | 1 | 0.349 |
| L × D | 4.222 | 3 | 0.238 | 3.944 | 3 | 0.268 |
| *Garden Slope Estimates* | Slope | SE |  | Slope | SE |  |
| Lafayette | – 0.251 | 0.115 |  | – 0.255 | 0.124 |  |
| Mississauga | 0.027 | 0.120 |  | 0.021 | 0.131 |  |
| Montpellier | 0.061 | 0.172 |  | 0.097 | 0.174 |  |
| Uppsala | – 0.099 | 0.113 |  | – 0.091 | 0.116 |  |
| Predictor | <u># of Flower Heads</u> |  |  | <u>Seed Set Mass</u> |  |  |
| | $\chi^2$ | df | P | $\chi^2$ | df | P |
| Continent of Origin (O) | 2.270 | 1 | 0.132 | 1.136 | 1 | 0.286 |
| Garden Location (L) | **247.413** | **3** | **< 0.001** | **39.550** | **3** | **< 0.001** |
| Climatic Distance (D) | **13.326** | **1** | **< 0.001** | **14.168** | **1** | **< 0.001** |
| O × L | 1.415 | 3 | 0.702 | 4.059 | 3 | 0.255 |
| O × D | 0.271 | 1 | 0.602 | 0.873 | 1 | 0.350 |
| L × D | **12.474** | **3** | **0.006** | **8.553** | **3** | **0.036** |
| *Garden Slope Estimates* | Slope | SE |  | Slope | SE |  |
| Lafayette | – 0.597 | 0.168 |  | – 0.674 | 0.223 |  |
| Mississauga | – 0.169 | 0.170 |  | – 0.273 | 0.217 |  |
| Montpellier | – 0.037 | 0.252 |  | 0.053 | 0.316 |  |
| Uppsala | 0.036 | 0.147 |  | 0.002 | 0.185 |  |
| Predictor | <u>Survival</u> |  |  | <u>Maximum Plant Area</u> |  |  |
| | $\chi^2$ | df | P | $\chi^2$ | df | P |
| Continent of Origin (O) | 2.408 | 1 | 0.121 | 0.924 | 1 | 0.336 |
| Garden Location (L) | **224.949** | **3** | **< 0.001** | **285.680** | **3** | **< 0.001** |
| Climatic Distance (D) | **6.633** | **1** | **0.010** | 0.582 | 1 | 0.446 |
| O × L | 6.049 | 3 | 0.109 | 6.467 | 3 | 0.091 |
| O × D | 0.093 | 1 | 0.761 | 1.752 | 1 | 0.186 |
| L × D | 4.609 | 3 | 0.203 | 5.620 | 3 | 0.132 |
| *Garden Slope Estimates* | Slope | SE |  | Slope | SE |  |
| Lafayette | – 0.109 | 0.111 |  | – 0.119 | 0.124 |  |
| Mississauga | – 0.010 | 0.117 |  | – 0.150 | 0.138 |  |
| Montpellier | 0.032 | 0.167 |  | 0.609 | 0.339 |  |
| Uppsala | – 0.260 | 0.100 |  | – 0.214 | 0.126 |  |

| Predictor | <u>Growth Rate</u> |  |  | <u>Herbivory</u> |  |  |
| --- | --- | --- | --- | --- | --- | --- |
| | $\chi^2$ | df | *P* | $\chi^2$ | df | *P* |
| Continent of Origin (O) | 0.094 | 1 | 0.760 | 0.088 | 1 | 0.766 |
| Garden Location (L) | **69.948** | **3** | **< 0.001** | **143.951** | **3** | **< 0.001** |
| Climatic Distance (D) | 0.087 | 1 | 0.768 | 1.070 | 1 | 0.301 |
| O × L | 1.207 | 3 | 0.751 | 6.613 | 3 | 0.085 |
| O × D | 0.049 | 1 | 0.825 | 0.010 | 1 | 0.920 |
| L × D | 2.873 | 3 | 0.412 | 2.226 | 3 | 0.527 |

| *Garden Slope Estimates* | Slope | SE | Slope | SE |
| --- | --- | --- | --- | --- |
| Lafayette | 0.001 | 0.003 | 0.177 | 0.155 |
| Mississauga | − 0.004 | 0.003 | 0.055 | 0.127 |
| Montpellier | 0.002 | 0.006 | − 0.358 | 0.346 |
| Uppsala | 0.002 | 0.002 | 0.079 | 0.136 |

Note: Significant effects ( $P < 0.05$ ) are listed in bold. Slope estimates and standard errors (SE) are listed individually for each common garden location.

### Q2: Does T. repens exhibit stronger local adaptation in its native range than in its introduced range?

We observed strong local adaptation in both the native and introduced ranges of *T. repens*, but the presence of local adaptation was dependent on where populations originated. This result is explained by a three-way interaction between continent of origin, continent of the common garden, and climatic distance, which affected all quantitative sexual fitness components, survival, and growth rate of *T. repens* (**Table 2**). Specifically, populations only exhibited strong signatures of local adaptation when they were transplanted to common gardens on the same continent from which they originated (**Figure 3**). For number of flower heads, in common gardens located in the native range (Europe), the slope for European *T. repens* populations was 12× steeper than the slope for North American populations. In common gardens located in the introduced range (North America), the slope for North American *T. repens* populations was 20× steeper than the slope for European populations. Similarly, for seed set mass, in common gardens located in Europe, the slope for European *T. repens* populations was 24× steeper than the slope for North American populations. In common gardens located in North America, the slope for North American *T. repens* populations was 7.8× steeper than the slope for European populations. The main effect of climatic distance affected whether plants flowered and produced seeds, but climatic distance did not interact with the continent of origin or the continent of the common garden to affect these fitness components (**Table 2**; **Figure S4**). However, climatic distance and continent of origin did interact to affect maximum plant area and herbivory. Together across all gardens, plants from European origins that were transplanted shorter climatic distances grew larger, despite experiencing greater levels of herbivory, while size and herbivory for plants from North American origins were not significantly affected by climatic distance overall (**Table 2**; **Figure S4**).

**Figure 3.**
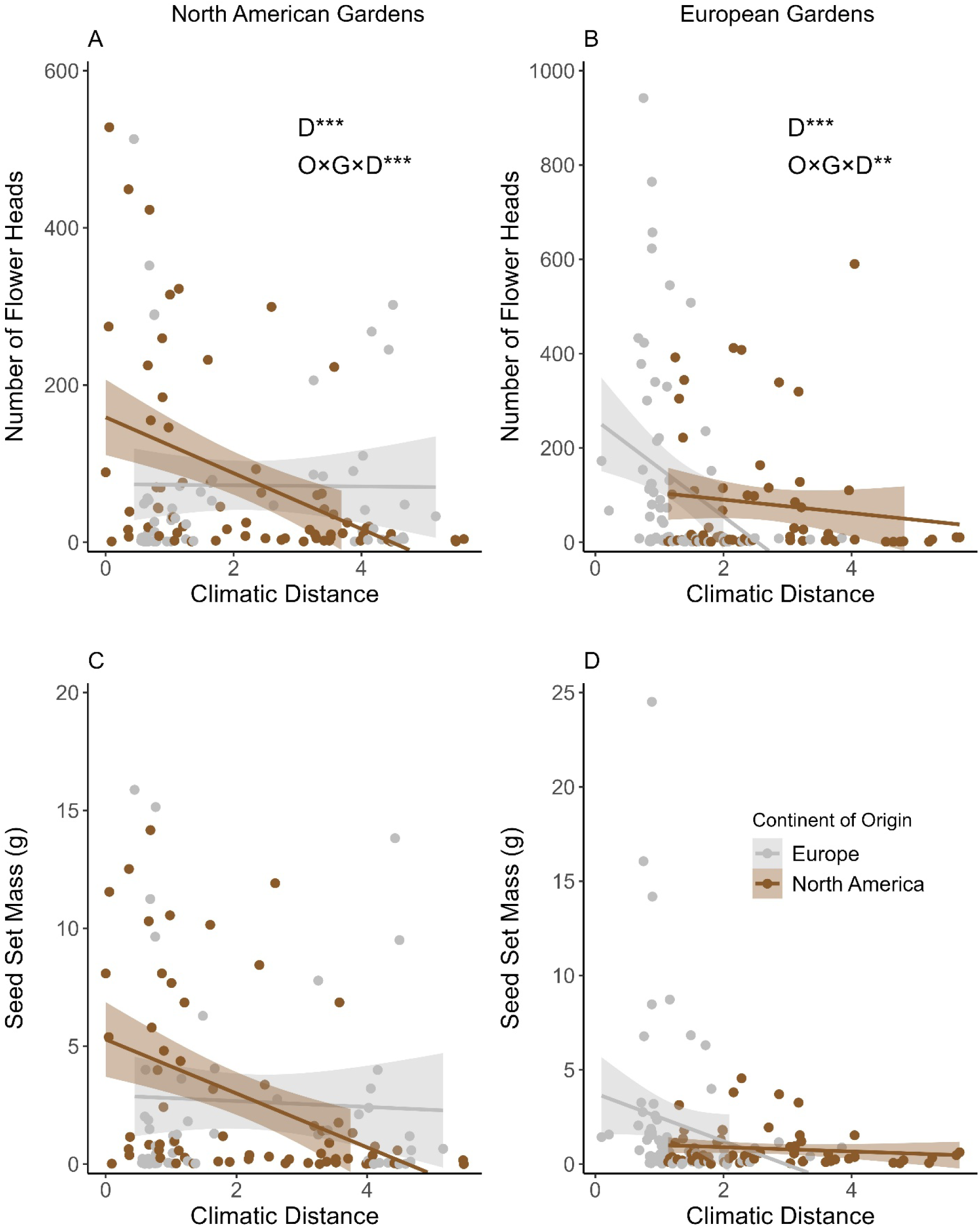
The effect of climatic distance on the mean number of flower heads produced by *T. repens* populations originating from its introduced (North America, orange) and native (Europe, pink) ranges and transplanted into common gardens in the (**A**) introduced and (**B**) native ranges. The effect of climatic distance on the mean seed set mass produced by *T. repens* populations originating from its introduced and native ranges and transplanted into common gardens in the (**C**) introduced and (**D**) native ranges. Shaded areas represent 95% confidence intervals around each line, while asterisks represent *P* values of < 0.05 for relevant fixed effects of Climatic Distance (D) and its interaction with Continent of Origin (O) and Continent of Garden (G).

**Table 2.** Results of mixed effects models assessing how continent of origin, continent of garden, and climatic distance affected the fitness of *T. repens*.

| Predictor | <u>Plants Producing</u><br><u>Flowers</u> |  | <u>Plants Producing</u><br><u>Seeds</u> |  |
| --- | --- | --- | --- | --- |
| | $\chi^2$ | *P* | $\chi^2$ | *P* |
| Continent of Origin (O) | 2.033 | 0.154 | 1.754 | 0.185 |
| Continent of Garden (G) | **9.184** | **0.002** | 1.295 | 0.255 |
| Climatic Distance (D) | **28.963** | **< 0.001** | **17.269** | **< 0.001** |
| O × G | 0.677 | 0.411 | 0.942 | 0.332 |
| O × D | 2.033 | 0.154 | 0.036 | 0.850 |
| G × D | 3.194 | 0.074 | 2.490 | 0.115 |
| O × G × D | 0.501 | 0.479 | 0.026 | 0.871 |
| *O × G Slope Estimates* | Slope | SE | Slope | SE |
| North America/North America | – 0.207 | 0.067 | – 0.241 | 0.076 |
| North America/Europe | – 0.072 | 0.096 | – 0.086 | 0.097 |
| Europe/North America | – 0.363 | 0.071 | – 0.231 | 0.077 |
| Europe/Europe | – 0.069 | 0.178 | – 0.038 | 0.181 |
| Predictor | <u># of Flower Heads</u> |  | <u>Seed Set Mass</u> |  |
| | $\chi^2$ | *P* | $\chi^2$ | *P* |
| Continent of Origin (O) | 0.514 | 0.474 | 0.028 | 0.867 |
| Continent of Garden (G) | 0.471 | 0.493 | 0.037 | 0.847 |
| Climatic Distance (D) | **14.310** | **< 0.001** | **16.190** | **< 0.001** |
| O × G | 0.957 | 0.328 | 0.481 | 0.488 |
| O × D | 1.214 | 0.271 | 2.991 | 0.084 |
| G × D | 1.412 | 0.235 | 0.258 | 0.612 |
| O × G × D | **19.603** | **< 0.001** | **9.728** | **0.002** |
| *O × G Slope Estimates* | Slope | SE | Slope | SE |
| North America/North America | – 0.482 | 0.125 | – 0.631 | 0.139 |
| North America/Europe | – 0.237 | 0.162 | – 0.179 | 0.169 |
| Europe/North America | 0.097 | 0.139 | – 0.006 | 0.151 |
| Europe/Europe | – 1.351 | 0.301 | – 0.826 | 0.313 |
| Predictor | <u>Survival</u> |  | <u>Maximum Plant</u><br><u>Area</u> |  |
| | $\chi^2$ | *P* | $\chi^2$ | *P* |
| Continent of Origin (O) | 0.106 | 0.745 | 0.797 | 0.372 |
| Continent of Garden (G) | **41.220** | **< 0.001** | **170.553** | **< 0.001** |
| Climatic Distance (D) | **10.236** | **0.001** | **58.341** | **< 0.001** |
| O × G | 0.000 | 0.993 | 2.375 | 0.123 |
| O × D | 1.534 | 0.216 | **34.173** | **< 0.001** |
| G × D | **13.870** | **< 0.001** | 1.784 | 0.182 |
| O × G × D | **6.515** | **0.011** | 0.678 | 0.410 |

| *O × G Slope Estimates* | Slope | SE | Slope | SE |
| --- | --- | --- | --- | --- |
| North America/North America | – 0.133 | 0.067 | – 0.109 | 0.078 |
| North America/Europe | 0.062 | 0.078 | – 0.005 | 0.122 |
| Europe/North America | – 0.357 | 0.069 | – 0.721 | 0.073 |
| Europe/Europe | 0.335 | 0.152 | – 0.385 | 0.230 |

|  | <u>Growth Rate</u> |  | <u>Herbivory</u> |  |
| --- | --- | --- | --- | --- |
| Predictor | $\chi^2$ | *P* | $\chi^2$ | *P* |
| Continent of Origin (O) | 0.053 | 0.817 | 0.196 | 0.658 |
| Continent of Garden (G) | 1.468 | 0.226 | **21.180** | **< 0.001** |
| Climatic Distance (D) | **18.137** | **< 0.001** | **15.810** | **< 0.001** |
| O × G | 0.465 | 0.495 | 0.669 | 0.413 |
| O × D | **5.145** | **0.023** | **23.686** | **< 0.001** |
| G × D | 2.263 | 0.133 | 0.240 | 0.624 |
| O × G × D | **7.812** | **0.005** | 2.178 | 0.140 |

| *O × G Slope Estimates* | Slope | SE | Slope | SE |
| --- | --- | --- | --- | --- |
| North America/North America | 0.002 | 0.002 | 0.069 | 0.091 |
| North America/Europe | 0.003 | 0.002 | – 0.122 | 0.135 |
| Europe/North America | 0.012 | 0.002 | – 0.591 | 0.094 |
| Europe/Europe | – 0.003 | 0.004 | – 0.311 | 0.257 |

Note: Significant effects ( $P < 0.05$ ) are listed in bold. All predictors are assessed using one degree of freedom. Slope estimates and standard errors (SE) are listed individually for each common garden location.

### Q3: What are the roles of cyanogenesis and herbivory in driving local adaptation across space?

Both *Ac/ac* and *Li/li* exhibited strong latitudinal gene frequency clines among *T. repens* populations from both continents (**Table S8**). However, the lack of an interaction between latitude and continent of origin for both loci indicates no differences in the steepness of cyanogenesis clines between the native and introduced ranges of *T. repens* (**Figure S5**). The main effect of cyanotype affected survival, whether plants produced flowers and/or seeds, and maximum plant area, with cyanogenic plants generally exhibiting greater fitness than acyanogenic plants, regardless of the environment in which the common garden was found (**Table S9**). However, cyanotype alone did not affect the level of herbivory occurring on *T. repens* plants, and cyanotype did not interact with either common garden location or climatic distance to affect any fitness components or herbivory (**Table S9**).

### Q4: What bioclimatic variables have the strongest impact on fitness across space in T. repens?

Climatic variables related to both temperature and precipitation predicted variation in fitness in *T. repens*, but the most influential predictors varied between garden locations, with patterns emerging based on latitude (**Tables S3-S6**, red highlights). Overall, 58% of the most influential predictors were temperature-based (mean annual temperature, maximum temperature of the warmest month, or minimum temperature of the coldest month), while 42% were precipitation-based (annual precipitation, precipitation of the wettest month, or precipitation of the driest month). In the more southern gardens on each continent (Lafayette and Montpellier), 67% of the most influential predictors were precipitation-based, while only 33% were temperature-based. Conversely, in the more northern gardens on each continent (Mississauga and Uppsala), 80% of the most influential predictors were temperature-based, while only 20% were precipitation-based. We did not find evidence that the native and introduced ranges of *T. repens* were differentially affected by temperature or precipitation variables. In the native range (Uppsala and Montpellier, combined), 57% of the most influential predictors were temperature-based and 43% were precipitation-based. In the introduced range (Mississauga and Lafayette, combined), 58% of the most influential predictors were temperature-based, while 42% were precipitation-based.

### Q5: Are there adaptation lags to climate change in T. repens?

North American populations of *T. repens* demonstrated evidence of an adaptation lag in the northern garden in the introduced range (**Table 3**). Survival, seed set mass, and maximum plant area all exhibited a significant quadratic relationship with ΔMAT between locations of origin and the Mississauga common garden (**Figure 4**). A quadratic relationship centred on a ΔMAT of zero would indicate local adaptation to the climate surrounding the common garden, while a relationship centred on a ΔMAT > 0 would represent an adaptation lag. For the proportion of plants survived per population, the peak of the quadratic relationship experienced a significant shift to a climate of origin with a MAT 3.4°C greater than Mississauga (*χ^2^_1,4_*= 4.159, *P* = 0.041, ΔAIC = −2.159), with the predicted maximum survival being 5.3% greater than at a ΔMAT of zero. For mean seed set mass per population, the peak of the quadratic relationship occurred at a MAT of 2.0°C greater than Mississauga, with the predicted maximum seed set mass being 5.2% greater than at a ΔMAT of zero. However, this shift was not statistically significant (*χ^2^_1,5_* = 0.640, *P* = 0.423, ΔAIC = 1.360). For mean maximum plant area per population, the peak of the quadratic relationship experienced a shift to a climate of origin with a MAT of 2.1°C (*χ^2^_1,5_* = 3.721, *P* = 0.054, ΔAIC = −1.721) greater than Mississauga, with the predicted maximum plant area being 3.9% greater than at a ΔMAT of zero. Along our transect of sampled populations, a shift in the peak of the quadratic relationship towards populations 2.0-3.1°C warmer than the Mississauga common garden represents a 2.1-3.2° shift in latitude towards the equator. Additionally, whether plants flowered or produced seeds increased linearly with ΔMAT, indicating plants of warmer origins were more likely to flower and produce seeds (**Figure 4**), while the number of flower heads and growth rate did not exhibit any significant linear or quadratic trends with ΔMAT (**Table 3**).

**Figure 4.**
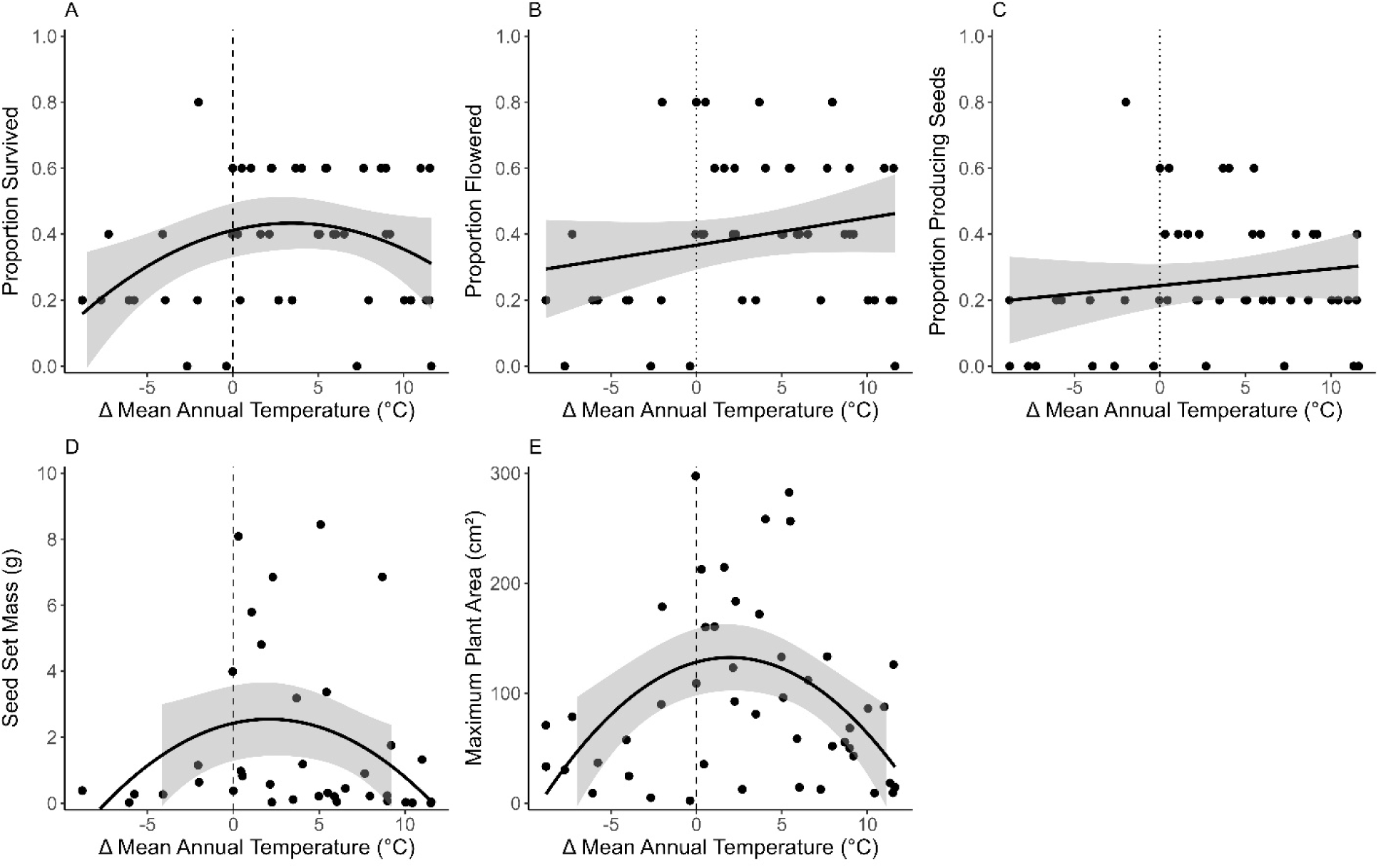
The effects of the change in mean annual temperature (ΔMAT) on fitness of North American populations of *T. repens* grown in the Mississauga common garden. (**A**) Proportion of plants survived, (**B**) proportion of plants that produced flowers, (**C**) proportion of plants that produced seeds, (**D**) mean seed set mass, and (**E**) mean maximum plant area per population all exhibited evidence of an adaptation lag in response to climate warming, indicated by a positive linear trend or a shift in the peak of the quadratic curve towards populations with warmer climates of origin (denoted as x-axis values > 0). Shaded areas represent 95% confidence intervals around each line.

**Table 3.** Results of mixed effects models assessing how change in mean annual temperature (ΔMAT) affected the fitness of North American *T. repens* populations in the Mississauga common garden (linear and quadratic fits).

| Predictor | <u>Plants Producing</u><br><u>Flowers</u> |  | <u>Plants Producing</u><br><u>Seeds</u> |  | <u># of Flower</u><br><u>Heads</u> |  | <u>Seed Set</u><br><u>Mass</u> |  |
| --- | --- | --- | --- | --- | --- | --- | --- | --- |
| | $\chi^2$ | *P* | $\chi^2$ | *P* | $\chi^2$ | *P* | $\chi^2$ | *P* |
| ΔMAT | **7.425** | **0.006** | **5.397** | **0.020** | 0.255 | 0.614 | 0.644 | 0.422 |
| ΔMAT<sup>2</sup> | **8.077** | **0.004** | **7.609** | **0.006** | 2.908 | 0.088 | **8.542** | **0.003** |

| Predictor | <u>Survival</u> |  | <u>Maximum Plant</u><br><u>Area</u> |  | <u>Growth Rate</u> |  |
| --- | --- | --- | --- | --- | --- | --- |
| | $\chi^2$ | *P* | $\chi^2$ | *P* | $\chi^2$ | *P* |
| ΔMAT | 3.758 | 0.053 | 3.799 | 0.051 | 0.002 | 0.962 |
| ΔMAT<sup>2</sup> | **4.313** | **0.038** | **6.741** | **0.009** | 1.808 | 0.179 |

Note: Significant effects ( $P < 0.05$ ) are listed in bold. All predictors are assessed using one degree of freedom.

## Discussion

The aim of our study was to provide insight into the potential for rapid local adaptation in introduced plant populations in response to novel environments across space and time. The results of our transcontinental experiment in the native (Europe) and introduced (North America) ranges of *T. repens* lead to two main conclusions: 1) rapid local adaptation to climatic variation across space has contributed to the ecological success of *T. repens* in its introduced range; 2) climate warming has resulted in some evidence of a weakening of local adaptation over time in the introduced range of *T. repens.* Our conclusions are based on five experimental results. First, *T. repens* exhibited local adaptation to climate overall, with greater fitness advantages for local populations at lower latitudes (Q1). Second, in European common gardens, only native European populations exhibited local adaptation, while in North American common gardens, only introduced North American populations exhibited local adaptation (Q2). Third, cyanogenesis did not differentially affect the fitness of *T. repens* in different environments (Q3). Fourth, both temperature- and precipitation-based bioclimatic variables predicted fitness, but temperature was a stronger determinant of fitness at higher latitudes, while precipitation was a stronger determinant of fitness at lower latitudes (Q4). Fifth, there is some evidence consistent with an adaptation lag in response to climate warming, but the signature was weak, and this pattern was only clear in the northern common garden of the introduced range (Q5). Below we discuss our results in the context of the role of rapid local adaptation in the success of non-native species and how introduced populations may (or may not) be able to adapt to future climate change.

### Strength of local adaptation is dependent on latitude

Spatial variation in biotic and abiotic factors across a species’ range could affect its ability to adapt to different environments within that range. We found that *T. repens* exhibited significant local adaptation to the abiotic environment based on a measure of climatic distance that encompassed mean annual temperature and precipitation along with annual temperature and precipitation extremes. However, local adaptation differed significantly in strength depending on the latitude of common gardens. Specifically, local adaptation was stronger at lower latitudes in both the native and introduced ranges of *T. repens* when fitness was measured as sexual reproductive success (i.e., number of flower heads and seed set mass). Our results are consistent with transplant experiments in other plant species, which found sexual fitness advantages for local versus foreign plants to be more pronounced at lower latitudes (Colautti and Barrett, 2013; Toräng et al., 2015), likely at least partially due to life history trade-offs across large latitudinal ranges (Li et al., 2015; Bastias et al., 2024). In fact, life history trade-offs in *T. repens* contribute to local adaptation across divergent environments, with southern populations investing more into sexual reproduction and northern populations investing more into vegetative growth (Wright et al., 2021). If warmer climates are generally more conducive to sexual reproduction than cooler climates, this may explain why sexual fitness advantages for local populations were more pronounced in Lafayette and Montpellier in our study, while climatic distance had more limited effects on vegetative growth.

### Local adaptation in native and introduced ranges

Non-native species could become more widespread and persistent by rapidly adapting to novel environments. For example, latitudinal cyanogenesis clines in introduced North American *T. repens* populations formed due to rapid local adaptation and sorting of existing genotypes post-introduction (Kooyers and Olsen, 2012; Kooyers and Olsen, 2013; Olsen et al., 2021). We found that populations of *T. repens* only exhibited strong signatures of local adaptation to climate when transplanted into common gardens within the same range as they originated. The observed strong local fitness advantages for North American populations within the North American gardens are therefore indicative of rapid local adaptation of *T. repens* within the past 400 years since its introduction. Strong selection on introduced populations provides opportunity for rapid local adaptation in non-native species on even shorter timescales than exhibited by *T. repens* (Colautti and Lau, 2015; Simón-Porcar et al., 2021). A meta-analysis by Oduor et al. (2016) found that non-native populations exhibited local adaptation just as frequently as native populations, and that the strength of local adaptation was not affected by time since introduction (ranging from as little as 20 to as many as 500 years). More specifically, previous studies on invasion in *Hypericum canariense* (Dlugosch and Parker, 2008), *Eschcholzia californica* (Leger and Rice, 2007), and *Lythrum salicaria* (Colautti and Barrett, 2013) indicate local adaptation in introduced ranges within 50, 150, and 200 years, respectively. Taken together, our results strengthen the mounting evidence that rapid local adaptation to climate plays an important role in the ecological success of non-native species in their new range.

Interestingly, *T. repens* populations from the introduced range did not exhibit signatures of local adaptation to similar climates when transplanted back into their native range. Similarly, native populations did not maintain their strong signatures of local adaptation to climate when transplanted into the introduced range. One important caveat is the overall narrower range of climatic distances among European populations (particularly within European common gardens), despite our widespread sampling (Figure S2), which could have limited our ability to compare the strength of local adaptation between ranges (Lucas et al., 2024). Additionally, while our common gardens are robust to detecting local adaptation caused by genotype × environment interactions (Bock et al., 2018), we also acknowledge that phenotypic and/or transgenerational plasticity (whose determination was beyond the scope of this study) may also be important for successful establishment or spread of non-native species (Hulme, 2008; Valladares et al., 2014; de Villemereuil et al., 2018). Despite these considerations, our findings demonstrate a clear cost of rapid adaptation, with local fitness advantages not extending across continents, even when accounting for a comprehensive measure of climatic distance between source environments and common garden environments on both continents. Explanations for this lack of maintenance of local adaptation across continents could be due to unmeasured biotic or abiotic aspects of the environment.

Abiotic explanations for the lack of local adaptation across continents could involve shifts in photoperiod or precipitation regimes between Europe and North America due to similar climates on the two continents being associated with different latitudes. In general, European environments have longer day lengths in summer and shorter day lengths in winter than those of similar climates in North America, with differences being more pronounced at progressively higher latitudes. This photoperiodic shift between the continents could have affected the initiation of flowering and/or the timing of peak flowering for populations that were transplanted across continents, leading to differences in the strength of local adaptation. Halbritter et al. (2015) found asymmetries in the strength of local adaptation in reciprocally-transplanted high-elevation and high-latitude *Plantago lanceolata* and *Plantago major* populations with similar climates of origin, indicating a potential role of photoperiod in local adaptation. Additionally, populations of an obligate long-day plant, *Mimulus guttatus*, exhibit clinal variation in the critical photoperiod required to initiate flowering, potentially indicating a role of photoperiod in driving local adaptation (Kooyers et al., 2015). Saikkonen et al. (2012) speculated that the transition to less extreme yearly variation in photoperiod at lower latitudes might facilitate greater ecological success of European species introduced to North America than vice versa, which could be especially true of a facultative long-day plant like *T. repens* (Beatty, 1956). However, our results do not provide evidence of increased success of European populations in North American gardens compared to that of North American populations in European gardens, indicating that other environmental factors are likely at play. Additionally, although we accounted for mean annual precipitation and yearly precipitation extremes in our study, these variables do not fully capture the exact timing and volume of precipitation, and the resulting hydrological environment experienced by *T. repens* plants in Europe versus North America. For example, in the Montpellier garden in southern Europe, plants will experience a Mediterranean climate that is not only drier but also more seasonally variable, while in the Lafayette garden in southern North America, plants will experience a sub-tropical environment that is wetter with more limited seasonal variation. This shift in precipitation availability has caused introduced *T. repens* populations to evolve greater drought avoidance as opposed to the more desiccation tolerant strategy observed in Mediterranean populations (Hendrickson et al., 2024).

While our experiment offers strong tests of local adaptation to both biotic and abiotic factors in the native and introduced ranges of a cosmopolitan plant, one limitation of this study is an inability to determine the relative strength of soil conditions compared to climate as drivers of local adaptation. Planting *T. repens* individuals from each population along with the natural soil and microbial communities from their location of origin was infeasible considering our study design and governmental regulations on importation of soil and bacteria. Instead, the best alternative was to maintain consistent treatment of all individuals in all gardens, with *T. repens* individuals planted into the natural soil at each common garden site. It is important to recognize that local edaphic factors have the potential to provide fitness advantages in *T. repens*, as has been demonstrated in other systems (Macel et al., 2007; Murray-Stoker and Johnson, 2024). Despite this, signatures of local adaptation to climate were still evident across multiple fitness components, strengthening our confidence in the observed patterns of rapid adaptation across ranges. However, a future study exploring the roles of edaphic and other abiotic factors could provide more comprehensive insight into how introduced plant populations adapt to new ranges.

### Cyanogenesis did not affect local adaptation in either the native or introduced range

The biotic environment could also partially explain the lack of local adaptation across continents based on climatic distance (Hargreaves et al, 2020). In line with classic hypotheses in invasion biology, herbivory could exhibit substantial differences across environments based on expected shifts in herbivore communities between ranges, which could facilitate success of a plant species post-introduction (Coley et al., 1985; Blossey and Nötzold, 1995; Keane and Crawley, 2002). If herbivory was likely to explain our results, we might expect to see signatures of local adaptation based on the presence of chemical defenses in *T. repens* plants. Among our sampled populations, we demonstrated rapid evolution of latitudinal clines in the presence of both *Ac* and *Li* alleles in the introduced range of *T. repens*, based on their similar strength to clines in the native range, demonstrating potential for involvement of cyanogenesis in local adaptation. However, our experimental results indicate that HCN did not confer higher fitness to local plants compared to foreign plants in any garden. In fact, local plants exhibited signatures of local adaptation despite experiencing similar or greater levels of herbivory than populations transplanted from further climatic distances. Additionally, experimental plants in the introduced range exhibited greater herbivory than plants in the native range, which is contrary to the prediction that *T. repens* was released from natural enemies in its native range (Keane and Crawley, 2002). Previous studies on local adaptation in *T. repens* have similarly found that HCN only had marginal, if any, effects on fitness (Wright et al., 2018; Wright et al., 2021) or have found weaker clines in the introduced range than the native range (Daday, 1954a; Daday, 1958; Innes et al., 2022). Taken together, our study contrasts classic invasion hypotheses, providing little evidence of a role of biotic factors (cyanogenesis or herbivory) in driving local adaptation of *T. repens* to its introduced range.

Our findings contrast with previous work that has speculated on the involvement of trade-offs in the costs and benefits of cyanogenesis or its metabolic components in adaptation to contrasting environments. For example, exacerbated ecological or allocation costs of producing HCN or its metabolic components in cooler climates could pair with exacerbated benefits of cyanogenesis in warmer climates with greater herbivore pressure (Daday, 1954a; Daday, 1965; Kooyers et al., 2018; Fadoul et al., 2023). In our study, cyanogenic plants often demonstrated a fitness advantage (i.e., benefits outweighing costs), even in cooler environments like Mississauga. However, despite these fitness advantages, we did not find evidence for HCN as a deterrent to herbivory in *T. repens*, indicating that greater prevalence of cyanogenesis in warmer climates is likely not based on an increased ability to defend against greater herbivore pressure. Previous work instead indicates the potential for cyanogenic glucosides and/or linamarase, individually, to be advantageous in periods of drought through the storage and recycling of nitrogen (Møller, 2010; Kooyers et al., 2014; Machingura et al., 2016). However, we also do not find evidence of the metabolic components of cyanogenesis affecting the fitness of *T. repens* or its adaptation to local climates. It is important to highlight that we were unable to harvest enough tissue to determine the cyanotype of most experimental plants due to concerns over effects of tissue harvest on plant fitness and/or mortality, so the strength of our conclusions with respect to the role of cyanogenesis in adaptation are limited by low sample sizes. Additionally, we do not account for other biotic interactions, such as belowground herbivory, soil biota, or competition, for their potential involvement in selection and local adaptation in *T. repens* (Blossey and Nötzold, 1995; Mitchell et al., 2006; Pal et al., 2020; Sheng et al., 2022; Murray-Stoker and Johnson, 2024). Further field studies are needed to more explicitly test a diverse array of biotic factors as drivers of local adaptation to climate across large environmental gradients, including direct tests of the costs and benefits of the most prominent defense mechanism in *T. repens*.

### Spatial variation in environmental drivers of local adaptation

While *T. repens* exhibits local adaptation in its native and introduced ranges based on an overall measure of climatic distance, populations may spatially vary in the individual bioclimatic variables responsible for local adaptation. Our results across both ranges demonstrate that at higher latitudes, where local adaptation is weaker, temperature-based variables are stronger determinants of fitness, while at lower latitudes, where local adaptation is stronger, precipitation-based variables are stronger determinants of fitness. Therefore, it is possible that *T. repens* adapts more readily to local precipitation regimes than local temperature regimes, resulting in stronger signatures of local adaptation in lower latitude common gardens, consistent between the native and introduced ranges. This result is consistent with Siepielski et al. (2017), which found both precipitation and potential evapotranspiration to be stronger predictors of selection than temperature across numerous taxa. Alternatively, our results could be affected by the specific environments chosen for the common gardens. In our study, the climates in Lafayette and Montpellier are among the warmest that occur in the range of *T. repens*, while representing two opposing extremes in terms of precipitation regime, and therefore could have been especially difficult to tolerate for populations transplanted from cooler climates with less extreme precipitation regimes. In turn, Uppsala was somewhat isolated, both geographically and climatically, from the sampled *T. repens* populations, which could have affected our ability to detect local adaptation (or maladaptation) in that environment. Lastly, it is possible that recent climate warming may have already disrupted the ability of local populations to outperform foreign populations, thereby impacting the strength of local adaptation in certain locations.

### Evidence of adaptation lags in response to climate warming

The ability of populations to adapt to recent rapid climate warming over time is an important mechanism by which species can mitigate the risk of local extinction and other downstream ecological consequences. Previous studies have demonstrated the incidence of adaptation lags, whereby populations originating from warmer climates would outperform populations with very similar climatic origins to the common garden location (Wilczek et al., 2014; Aguirre-Liguori et al., 2019; Anderson and Wadgymar, 2020). These studies indicate the potential for a disruption in local adaptation, and even the possibility of maladaptation, due to climate warming that is occurring faster than local populations are able to adapt.

In our northern common garden in North America (Mississauga), populations with warmer origins (2-3°C in MAT) outperformed local populations in survival and biomass production (i.e., a proxy for vegetative reproduction) but not sexual fitness components. This finding may signal the presence of an adaptation lag and may also provide a partial explanation for the observed weaker local adaptation in Mississauga compared to Lafayette. However, the generalizability of our results should be considered carefully for a few reasons. First, contrary to our expectations, we provide no evidence of either local adaptation or maladaptation in Uppsala. In a large-scale common garden experiment with European populations of *Arabidopsis thaliana*, Wilczek et al. (2014) found that adaptation lags were more pronounced in higher latitude common garden locations, which was not the case in our study. It would also be pertinent to be able to test for spatial lags in the native range of *T. repens* to which we could directly compare spatial lags in the introduced range. Second, we tested for the presence of adaptation lags based solely on MAT, and Mississauga was the common garden location where MAT most consistently affected plant fitness compared to other bioclimatic variables. Fitness in lower latitude common gardens was instead more heavily influenced by precipitation-based variables, which may not be changing as consistently as temperature-based variables due to anthropogenic activity but could still play a significant role in maladaptation in response to changing environments (Nagy et al., 2024). It is therefore possible that adaptation lags to changing environments may not occur consistently across different geographic locations, based on the specific biotic and abiotic factors that most strongly impact local adaptation in those environments. Third, the MAT at the location of the Mississauga common garden in 2020 and 2021 was 1.6°C and 2.0°C warmer, respectively, than the average from the historical time period (1970-2000) used to calculate climatic distances in this study (data extracted from ClimateNA; Wang et al., 2016). Therefore, it may not be unexpected to see populations from warmer origins outperform local populations. The adaptation lag for survival does extend to a peak 3.4°C warmer than Mississauga, and the proportion of plants producing flowers and/or seeds increased linearly with ΔMAT, which still provides evidence for warm-adapted populations having an advantage that exceeds expectations based on the MAT during the years of the experiment. However, across our entire study, we are unable to make broad conclusions about future vulnerability of populations based on the rate of environmental change exceeding the rate of adaptation. Instead, we conclude that the fine-tuned nature of local adaptation (in terms of variation in strength, prominent driving factors, and the potential for maladaptation) to climate in *T. repens* highlights the multifaceted nature of rapid evolution in response to changing environments.

## Conclusions

Our study provides insight into the ability of introduced populations of a widespread plant to adapt to novel environments across space and over time. Signatures of local adaptation were only evident when populations were transplanted within the same range as they originated, indicating both strong and rapid local adaptation to the introduced range of *T. repens* in less than 400 years since introduction. Interestingly, the strength of local adaptation and the most important environmental predictors of plant fitness differed in the north and south of both the native and introduced range. Specifically, temperature was the most important driver of local adaptation in northern gardens, whereas precipitation was most important in southern gardens. This result indicates that there may be predictable drivers of local adaptation of non-native populations to novel environments, and that these drivers may vary along broad latitudinal gradients. Despite evidence of rapid adaptation to changing environments across space, the northern garden in the introduced range also provided some evidence of local maladaptation compared to populations with warmer origins, which is characteristic of an adaptation lag. Therefore, while local adaptation has occurred quickly in introduced populations of *T. repens* across space, a lack of rapid adaptation to environmental change over time could eventually result in severe erosion of fitness advantages for local populations in the future. Continuing to further our understanding of the ability of native and introduced populations to rapidly adapt to anthropogenic environmental change in space and time is vital to the integrity and maintenance of biodiversity in the face of future shifts in environment.

## Acknowledgements

We thank James Santangelo for assistance with initial seed collection, and Happy Gill, Jashandeep Nijjar, and Ruth Thomas for assisting with the generation of F1 plants for use in the experiment. We are indebted to Aaron Albano, Barry Albano, Paula Albano, Sarah Albano, Andre Daugeureaux, Hayden Fargo, Axel Hjelm, Antoniette Jansen, Eunice Karinho Betancourt, Erik Karlsson, Andreas Lundgren, Kavya Manikonda, Lindsay Miles, David Murray-Stoker, Bernabé Ramirez, Brittany Sutherland, Olivia Toth, Andrea Turcu, Mariona Vinyeta Medina, and Vicki Zhang for assistance in the field. Keerath Bhachu, Elana Maria, Zain Nasrullah, Nial Navaratne, Narindra Persaud, and Isabellio Vessio assisted with sample processing. We also thank the Louisiana Optical Network (LONI) for providing computation resources, along with Kris Horvath, Joan Lee, Kent Moore, Diane Ross, Vera Velasco, the technical platform “Plateforme des Terrains d’Expérience” (CEFE-CNRS), and the University of Louisiana Ecology Center for facilitating and supporting the set-up and execution of the experiment. This research was supported by an NSERC Discovery Grant, Canada Research Chair, and NSERC EWR Steacie Fellowship to MTJJ, a CNRS-University of Toronto Ph.D. Student Travel Grant to MTJJ and CV, NSF grants OIA-1920858 and DBI-2244712 to NJK, and an NSERC CGS Doctoral Award to LJA.

---

**Table S1.** Historical bioclimatic data (1970-2000) for each common garden site.

| Bioclimatic Variable | <u>Mississauga</u> | <u>Lafayette</u> | <u>Uppsala</u> | <u>Montpellier</u> |
| --- | --- | --- | --- | --- |
| BIO1 (°C) | 8.4 | 19.7 | 5.9 | 14.7 |
| BIO5 (°C) | 26.9 | 33.1 | 21.9 | 28.8 |
| BIO6 (°C) | − 9.1 | 4.4 | − 7.0 | 3.1 |
| BIO12 (mm) | 813 | 1578 | 568 | 657 |
| BIO13 (mm) | 79 | 158 | 73 | 97 |
| BIO14 (mm) | 49 | 110 | 26 | 21 |

Note: Bioclimatic variables are mean annual temperature (BIO1), maximum temperature of the warmest month (BIO5), minimum temperature of the coldest month (BIO6), annual precipitation (BIO12), precipitation of the wettest month (BIO13), and precipitation of the driest month (BIO14). Data was extracted from WorldClim (Version 2.1, February 2023).

**Table S2.** Bivariate Pearson’s correlation coefficients between all bioclimatic variables of interest, latitude, and climatic distance for each common garden.

|  | BIO1 | BIO5 | BIO6 | BIO12 | BIO13 | BIO14 | Latitude |
| --- | --- | --- | --- | --- | --- | --- | --- |
| BIO1 | — | — | — | — | — | — | — |
| BIO5 | 0.751 | — | — | — | — | — | — |
| BIO6 | 0.816 | 0.264 | — | — | — | — | — |
| BIO12 | 0.556 | 0.577 | 0.181 | — | — | — | — |
| BIO13 | 0.549 | 0.564 | 0.176 | 0.961 | — | — | — |
| BIO14 | 0.508 | 0.506 | 0.182 | 0.936 | 0.841 | — | — |
| Latitude | − 0.759 | − 0.937 | − 0.273 | − 0.733 | − 0.722 | − 0.647 | — |
| CD (Lafayette) | − 0.828 | − 0.754 | − 0.478 | − 0.921 | − 0.891 | − 0.874 | 0.851 |
| CD (Mississauga) | 0.661 | 0.728 | 0.235 | 0.789 | 0.824 | 0.693 | − 0.752 |
| CD (Montpellier) | 0.282 | 0.505 | − 0.162 | 0.741 | 0.761 | 0.676 | − 0.554 |
| CD (Uppsala) | 0.768 | 0.797 | 0.342 | 0.915 | 0.912 | 0.834 | − 0.882 |

Note: Bioclimatic variables are mean annual temperature (BIO1), maximum temperature of the warmest month (BIO5), minimum temperature of the coldest month (BIO6), annual precipitation (BIO12), precipitation of the wettest month (BIO13), and precipitation of the driest month (BIO14) for each of the 96 locations of source populations of *T. repens*. Climatic distances (CD) are included separately for the four common garden sites (Lafayette, Mississauga, Montpellier, and Uppsala). Bioclimatic data was extracted from WorldClim (Version 2.1, February 2023).

**Table S3.**
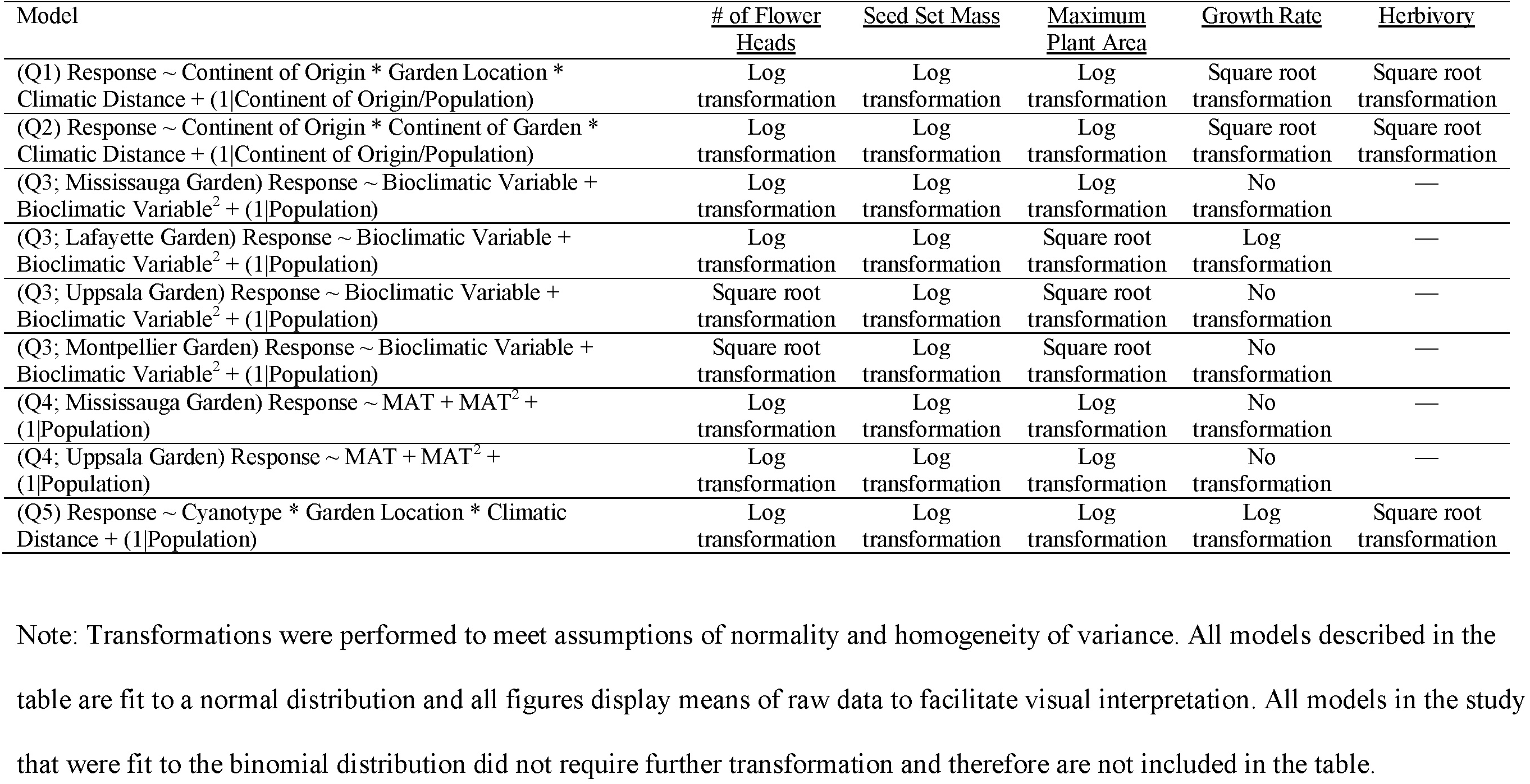
Summary of transformations used for each quantitative response variable.

**Table S4.** Sample sizes for each model used in the study, divided by the research question that each model set addresses and each of the eight response variables analyzed.

|  | <u>Survival</u> | <u>Plants</u><br><u>Producing</u><br><u>Flowers</u> | <u>Plants</u><br><u>Producing</u><br><u>Seeds</u> | <u># of Flower</u><br><u>Heads</u> | <u>Seed Set</u><br><u>Mass</u> | <u>Maximum</u><br><u>Plant Area</u> | <u>Growth Rate</u> | <u>Herbivory</u> |
| --- | --- | --- | --- | --- | --- | --- | --- | --- |
| Q1 | 1858 | 1858 | 1858 | 406 | 389 | 1262 | 730 | 949 |
| Q2 | 1858 | 1858 | 1858 | 406 | 389 | 1262 | 730 | 949 |
| Q3 | 554 | 554 | 554 | 218 | 211 | 553 | 343 | 507 |
| Q4 (Lafayette) | 465 | 465 | 465 | 77 | 66 | 323 | 129 | — |
| Q4 (Mississauga) | 465 | 465 | 465 | 117 | 113 | 451 | 188 | — |
| Q4 (Montpellier) | 470 | 470 | 470 | 102 | 102 | 127 | 102 | — |
| Q4 (Uppsala) | 458 | 458 | 458 | 110 | 108 | 361 | 311 | — |
| Q5 (Mississauga) | 235 | 235 | 235 | 64 | 61 | 229 | 91 | — |
| Q5 (Uppsala) | 233 | 233 | 233 | 58 | 57 | 179 | 153 | — |

**Table S5.** Results of mixed effects models assessing how individual bioclimatic variables (both linearly and quadratically) affected the fitness of *T. repens* in the Mississauga common garden.

| Predictor | <u>Survival</u> |  | <u>Plants Producing</u><br><u>Flowers</u> |  | <u>Plants Producing</u><br><u>Seeds</u> |  | <u># of Flower</u><br><u>Heads</u> |  | <u>Seed Set</u><br><u>Mass</u> |  | <u>Maximum</u><br><u>Plant Area</u> |  | <u>Growth Rate</u> |  |
| --- | --- | --- | --- | --- | --- | --- | --- | --- | --- | --- | --- | --- | --- | --- |
| | $\chi^2$ | *P* | $\chi^2$ | *P* | $\chi^2$ | *P* | $\chi^2$ | *P* | $\chi^2$ | *P* | $\chi^2$ | *P* | $\chi^2$ | *P* |
| BIO1 | **3.603** | **0.058** | **6.119** | **0.013** | **4.762** | **0.029** | 0.116 | 0.733 | 1.042 | 0.307 | 2.385 | 0.122 | 0.054 | 0.816 |
| BIO1<sup>2</sup> | **3.271** | **0.071** | **3.921** | **0.048** | **3.488** | **0.062** | 1.807 | 0.179 | **7.080** | **0.008** | **3.734** | **0.053** | 2.648 | 0.104 |
| BIO5 | 1.354 | 0.245 | **6.704** | **0.010** | **6.684** | **0.010** | 0.303 | 0.582 | 0.145 | 0.704 | 0.652 | 0.419 | 1.699 | 0.193 |
| BIO5<sup>2</sup> | 0.639 | 0.424 | **2.920** | **0.088** | 2.701 | 0.100 | 0.024 | 0.878 | 0.742 | 0.389 | 0.944 | 0.331 | 0.154 | 0.695 |
| BIO6 | 1.211 | 0.271 | 1.588 | 0.208 | 1.273 | 0.259 | 0.010 | 0.920 | 0.037 | 0.847 | 0.405 | 0.525 | 0.314 | 0.575 |
| BIO6<sup>2</sup> | 2.452 | 0.117 | **4.997** | **0.025** | **5.763** | **0.016** | 0.720 | 0.396 | **3.318** | **0.069** | 2.605 | 0.107 | 2.661 | 0.103 |
| BIO12 | 1.077 | 0.299 | 1.162 | 0.281 | 0.066 | 0.797 | 0.087 | 0.768 | 0.200 | 0.655 | 1.554 | 0.213 | 2.144 | 0.143 |
| BIO12<sup>2</sup> | 0.883 | 0.347 | 0.321 | 0.571 | 0.004 | 0.948 | 0.558 | 0.455 | **3.110** | **0.078** | 1.930 | 0.165 | **3.496** | **0.062** |
| BIO13 | 0.610 | 0.435 | 0.403 | 0.525 | 0.099 | 0.753 | 0.100 | 0.752 | 0.290 | 0.590 | 2.254 | 0.133 | **6.168** | **0.013** |
| BIO13<sup>2</sup> | 0.377 | 0.539 | 0.043 | 0.837 | 0.015 | 0.902 | 0.055 | 0.815 | 1.450 | 0.229 | 1.963 | 0.161 | **6.641** | **0.010** |
| BIO14 | 0.173 | 0.677 | 0.562 | 0.453 | 0.064 | 0.800 | 2.066 | 0.151 | 1.708 | 0.191 | 0.031 | 0.860 | 0.051 | 0.822 |
| BIO14<sup>2</sup> | 1.175 | 0.279 | 0.692 | 0.406 | 0.002 | 0.964 | 0.896 | 0.344 | **3.996** | **0.046** | 1.290 | 0.256 | 2.330 | 0.127 |

Note: Bioclimatic variables are mean annual temperature (BIO1), maximum temperature of the warmest month (BIO5), minimum temperature of the coldest month (BIO6), annual precipitation (BIO12), precipitation of the wettest month (BIO13), and precipitation of the driest month (BIO14). All variables were z-score standardized to allow for simultaneous direct comparisons of the relative strength of linear and quadratic predictor variables. Effects with $P < 0.1$ are bolded and the effect in red exhibited the lowest $P$ -value among all independent variables tested for a given fitness measure (among $P$ values $< 0.1$ ). All predictors are assessed using one degree of freedom.

**Table S6.** Results of mixed effects models assessing how individual bioclimatic variables (both linearly and quadratically) affected the fitness of *T. repens* in the Lafayette common garden.

| Predictor | <u>Survival</u> |  | <u>Plants Producing Flowers</u> |  | <u>Plants Producing Seeds</u> |  | <u># of Flower Heads</u> |  | <u>Seed Set Mass</u> |  | <u>Maximum Plant Area</u> |  | <u>Growth Rate</u> |  |
| --- | --- | --- | --- | --- | --- | --- | --- | --- | --- | --- | --- | --- | --- | --- |
| | $\chi^2$ | *P* | $\chi^2$ | *P* | $\chi^2$ | *P* | $\chi^2$ | *P* | $\chi^2$ | *P* | $\chi^2$ | *P* | $\chi^2$ | *P* |
| BIO1 | **3.562** | **0.059** | **4.562** | **0.033** | **4.498** | **0.034** | 0.514 | 0.474 | 2.559 | 0.110 | 0.006 | 0.940 | 1.062 | 0.303 |
| BIO1<sup>2</sup> | 2.237 | 0.135 | 0.767 | 0.381 | 0.556 | 0.456 | 0.291 | 0.590 | 0.051 | 0.821 | 0.000 | 0.988 | 1.467 | 0.226 |
| BIO5 | 0.109 | 0.741 | 0.866 | 0.352 | 0.421 | 0.516 | 0.112 | 0.738 | 1.950 | 0.163 | **6.484** | **0.011** | 0.005 | 0.946 |
| BIO5<sup>2</sup> | 0.491 | 0.484 | 0.012 | 0.912 | 0.108 | 0.742 | 0.032 | 0.859 | 0.426 | 0.514 | **8.242** | **0.004** | 0.047 | 0.829 |
| BIO6 | 2.680 | 0.102 | 1.279 | 0.258 | 2.156 | 0.142 | 1.672 | 0.196 | 1.100 | 0.294 | 0.052 | 0.820 | 1.252 | 0.263 |
| BIO6<sup>2</sup> | 2.375 | 0.123 | 0.264 | 0.607 | 0.473 | 0.492 | 0.009 | 0.923 | 0.000 | 0.991 | 0.013 | 0.909 | 1.746 | 0.186 |
| BIO12 | 0.307 | 0.580 | 2.493 | 0.114 | **3.151** | **0.076** | **2.989** | **0.084** | **7.048** | **0.008** | 0.426 | 0.514 | 0.046 | 0.830 |
| BIO12<sup>2</sup> | 0.006 | 0.937 | 0.399 | 0.528 | 0.659 | 0.417 | 0.910 | 0.340 | **2.831** | **0.092** | 0.955 | 0.328 | 0.024 | 0.877 |
| BIO13 | 1.157 | 0.282 | **5.384** | **0.020** | **5.918** | **0.015** | 2.139 | 0.144 | **5.549** | **0.018** | 0.171 | 0.679 | 0.745 | 0.388 |
| BIO13<sup>2</sup> | 0.169 | 0.681 | 1.328 | 0.249 | 1.640 | 0.200 | 0.231 | 0.631 | 1.577 | 0.209 | 0.781 | 0.377 | 0.591 | 0.442 |
| BIO14 | 0.377 | 0.539 | **4.254** | **0.039** | **5.182** | **0.023** | 1.860 | 0.172 | **3.346** | **0.067** | 0.879 | 0.349 | 0.404 | 0.525 |
| BIO14<sup>2</sup> | 0.047 | 0.829 | 1.485 | 0.223 | 1.823 | 0.177 | 0.551 | 0.458 | 0.955 | 0.328 | 1.407 | 0.236 | 0.442 | 0.506 |

Note: Bioclimatic variables are mean annual temperature (BIO1), maximum temperature of the warmest month (BIO5), minimum temperature of the coldest month (BIO6), annual precipitation (BIO12), precipitation of the wettest month (BIO13), and precipitation of the driest month (BIO14). All variables were z-score standardized to allow for simultaneous direct comparisons of the relative strength of linear and quadratic predictor variables. Effects with $P < 0.1$ are bolded and the effect in red exhibited the lowest $P$ -value among all independent variables tested for a given fitness measure (among $P$ values $< 0.1$ ). All predictors are assessed using one degree of freedom.

**Table S7.** Results of mixed effects models assessing how individual bioclimatic variables (both linearly and quadratically) affected the fitness of *T. repens* in the Uppsala common garden.

| Predictor | <u>Survival</u> |  | <u>Plants Producing</u><br><u>Flowers</u> |  | <u>Plants Producing</u><br><u>Seeds</u> |  | <u># of Flower</u><br><u>Heads</u> |  | <u>Seed Set</u><br><u>Mass</u> |  | <u>Maximum</u><br><u>Plant Area</u> |  | <u>Growth Rate</u> |  |
| --- | --- | --- | --- | --- | --- | --- | --- | --- | --- | --- | --- | --- | --- | --- |
| | $\chi^2$ | *P* | $\chi^2$ | *P* | $\chi^2$ | *P* | $\chi^2$ | *P* | $\chi^2$ | *P* | $\chi^2$ | *P* | $\chi^2$ | *P* |
| BIO1 | 1.749 | 0.186 | 0.332 | 0.564 | 0.226 | 0.635 | 0.295 | 0.587 | 0.004 | 0.948 | 0.512 | 0.475 | 0.209 | 0.648 |
| BIO1<sup>2</sup> | 0.114 | 0.736 | 1.291 | 0.256 | 1.137 | 0.286 | 0.583 | 0.445 | 0.009 | 0.923 | 0.242 | 0.622 | 1.389 | 0.239 |
| BIO5 | **3.906** | **0.048** | **5.261** | **0.022** | **5.104** | **0.024** | 0.377 | 0.539 | 1.694 | 0.193 | 0.083 | 0.773 | **4.633** | **0.031** |
| BIO5<sup>2</sup> | **6.392** | **0.011** | **6.781** | **0.009** | **6.643** | **0.010** | 0.361 | 0.548 | 1.665 | 0.197 | 0.072 | 0.789 | **5.401** | **0.020** |
| BIO6 | **9.121** | **0.003** | 0.457 | 0.499 | 0.532 | 0.466 | 0.048 | 0.826 | 0.325 | 0.569 | 0.656 | 0.418 | 0.717 | 0.397 |
| BIO6<sup>2</sup> | 0.722 | 0.396 | 2.530 | 0.112 | 2.381 | 0.123 | 0.779 | 0.377 | 0.027 | 0.870 | 0.001 | 0.979 | 0.841 | 0.359 |
| BIO12 | **9.972** | **0.002** | 0.778 | 0.378 | 0.782 | 0.377 | 0.973 | 0.324 | 0.001 | 0.972 | 0.535 | 0.465 | 0.454 | 0.501 |
| BIO12<sup>2</sup> | **12.362** | **< 0.001** | 1.197 | 0.274 | 1.269 | 0.260 | 0.862 | 0.353 | 0.005 | 0.944 | 0.536 | 0.464 | 1.929 | 0.165 |
| BIO13 | **8.552** | **0.003** | 0.603 | 0.437 | 0.713 | 0.398 | 0.519 | 0.471 | 0.021 | 0.886 | 0.379 | 0.538 | 0.075 | 0.785 |
| BIO13<sup>2</sup> | **11.256** | **< 0.001** | 1.104 | 0.293 | 1.343 | 0.247 | 0.516 | 0.473 | 0.051 | 0.822 | 0.240 | 0.624 | 1.219 | 0.270 |
| BIO14 | **6.626** | **0.010** | 1.180 | 0.277 | 1.058 | 0.304 | 2.045 | 0.153 | 0.177 | 0.674 | 0.742 | 0.389 | 0.515 | 0.473 |
| BIO14<sup>2</sup> | **8.578** | **0.003** | 1.696 | 0.193 | 1.673 | 0.196 | 1.797 | 0.180 | 0.001 | 0.981 | 1.155 | 0.283 | 1.931 | 0.165 |

Note: Bioclimatic variables are mean annual temperature (BIO1), maximum temperature of the warmest month (BIO5), minimum temperature of the coldest month (BIO6), annual precipitation (BIO12), precipitation of the wettest month (BIO13), and precipitation of the driest month (BIO14). All variables were z-score standardized to allow for simultaneous direct comparisons of the relative strength of linear and quadratic predictor variables. Effects with $P < 0.1$ are bolded and the effect in red exhibited the lowest $P$ -value among all independent variables tested for a given fitness measure (among $P$ values $< 0.1$ ). All predictors are assessed using one degree of freedom.

**Table S8.** Results of mixed effects models assessing how individual bioclimatic variables (both linearly and quadratically) affected the fitness of *T. repens* in the Montpellier common garden.

| Predictor | <u>Survival</u> |  | <u>Plants Producing</u><br><u>Flowers</u> |  | <u>Plants Producing</u><br><u>Seeds</u> |  | <u># of Flower</u><br><u>Heads</u> |  | <u>Seed Set</u><br><u>Mass</u> |  | <u>Maximum</u><br><u>Plant Area</u> |  | <u>Growth Rate</u> |  |
| --- | --- | --- | --- | --- | --- | --- | --- | --- | --- | --- | --- | --- | --- | --- |
| | $\chi^2$ | *P* | $\chi^2$ | *P* | $\chi^2$ | *P* | $\chi^2$ | *P* | $\chi^2$ | *P* | $\chi^2$ | *P* | $\chi^2$ | *P* |
| BIO1 | 0.248 | 0.618 | 1.006 | 0.316 | 1.024 | 0.312 | 1.599 | 0.206 | 1.397 | 0.237 | **3.389** | **0.066** | 1.002 | 0.317 |
| BIO1<sup>2</sup> | 0.317 | 0.574 | 0.027 | 0.869 | 0.089 | 0.765 | 1.771 | 0.183 | 1.210 | 0.271 | 0.487 | 0.485 | 0.001 | 0.974 |
| BIO5 | 0.076 | 0.782 | 0.360 | 0.548 | 0.263 | 0.608 | 0.660 | 0.417 | 0.043 | 0.836 | 1.633 | 0.201 | 0.017 | 0.898 |
| BIO5<sup>2</sup> | 0.793 | 0.373 | 0.780 | 0.377 | 1.250 | 0.264 | 0.024 | 0.877 | 1.274 | 0.259 | 0.357 | 0.550 | 0.376 | 0.540 |
| BIO6 | 0.519 | 0.471 | 0.688 | 0.407 | 0.685 | 0.408 | 1.039 | 0.308 | 2.139 | 0.144 | **6.489** | **0.011** | **5.509** | **0.019** |
| BIO6<sup>2</sup> | 0.816 | 0.366 | 0.225 | 0.635 | 0.287 | 0.592 | 0.130 | 0.719 | 0.070 | 0.791 | **3.978** | **0.046** | **2.862** | **0.091** |
| BIO12 | 0.202 | 0.653 | 0.244 | 0.621 | 0.094 | 0.760 | **2.797** | **0.094** | 1.208 | 0.272 | 2.606 | 0.106 | 0.459 | 0.498 |
| BIO12<sup>2</sup> | 0.151 | 0.697 | 0.513 | 0.474 | 0.430 | 0.512 | 2.577 | 0.108 | 0.586 | 0.444 | **5.722** | **0.017** | 1.085 | 0.298 |
| BIO13 | 0.715 | 0.398 | 1.007 | 0.316 | 0.602 | 0.438 | **3.470** | **0.063** | 1.323 | 0.250 | 0.701 | 0.403 | 0.071 | 0.790 |
| BIO13<sup>2</sup> | 0.582 | 0.446 | 1.204 | 0.273 | 0.960 | 0.327 | **3.588** | **0.058** | 1.240 | 0.266 | 1.344 | 0.246 | 0.000 | 0.987 |
| BIO14 | 0.000 | 0.991 | 0.021 | 0.884 | 0.065 | 0.799 | 2.539 | 0.111 | 2.177 | 0.140 | **2.758** | **0.097** | 1.317 | 0.251 |
| BIO14<sup>2</sup> | 0.502 | 0.579 | 0.895 | 0.344 | 1.017 | 0.313 | 1.880 | 0.170 | 0.515 | 0.473 | **7.215** | **0.007** | **3.214** | **0.073** |

Note: Bioclimatic variables are mean annual temperature (BIO1), maximum temperature of the warmest month (BIO5), minimum temperature of the coldest month (BIO6), annual precipitation (BIO12), precipitation of the wettest month (BIO13), and precipitation of the driest month (BIO14). All variables were z-score standardized to allow for simultaneous direct comparisons of the relative strength of linear and quadratic predictor variables. Effects with $P < 0.1$ are bolded and the effect in red exhibited the lowest $P$ -value among all independent variables tested for a given fitness measure (among $P$ values $< 0.1$ ). All predictors are assessed using one degree of freedom.

**Table S9.** Results of mixed effects models assessing how change in mean annual temperature (not z-score standardized) affected the fitness of European *T. repens* populations in the Uppsala common garden (linear and quadratic fits).

| Predictor | <u>Survival</u> |  | <u>Plants Producing</u><br><u>Flowers</u> |  | <u>Plants Producing</u><br><u>Seeds</u> |  | <u># of Flower</u><br><u>Heads</u> |  | <u>Seed Set</u><br><u>Mass</u> |  | <u>Maximum</u><br><u>Plant Area</u> |  | <u>Growth Rate</u> |  |
| --- | --- | --- | --- | --- | --- | --- | --- | --- | --- | --- | --- | --- | --- | --- |
| | $\chi^2$ | *P* | $\chi^2$ | *P* | $\chi^2$ | *P* | $\chi^2$ | *P* | $\chi^2$ | *P* | $\chi^2$ | *P* | $\chi^2$ | *P* |
| $\Delta\text{MAT}$ | 1.612 | 0.204 | 0.429 | 0.513 | 0.306 | 0.580 | 0.003 | 0.957 | 0.006 | 0.941 | 0.000 | 0.999 | 0.101 | 0.751 |
| $\Delta\text{MAT}^2$ | 2.267 | 0.132 | 0.595 | 0.440 | 0.440 | 0.507 | 0.028 | 0.867 | 0.070 | 0.792 | 0.004 | 0.953 | 0.184 | 0.668 |

Note: All predictors are assessed using one degree of freedom.

**Table S10.**
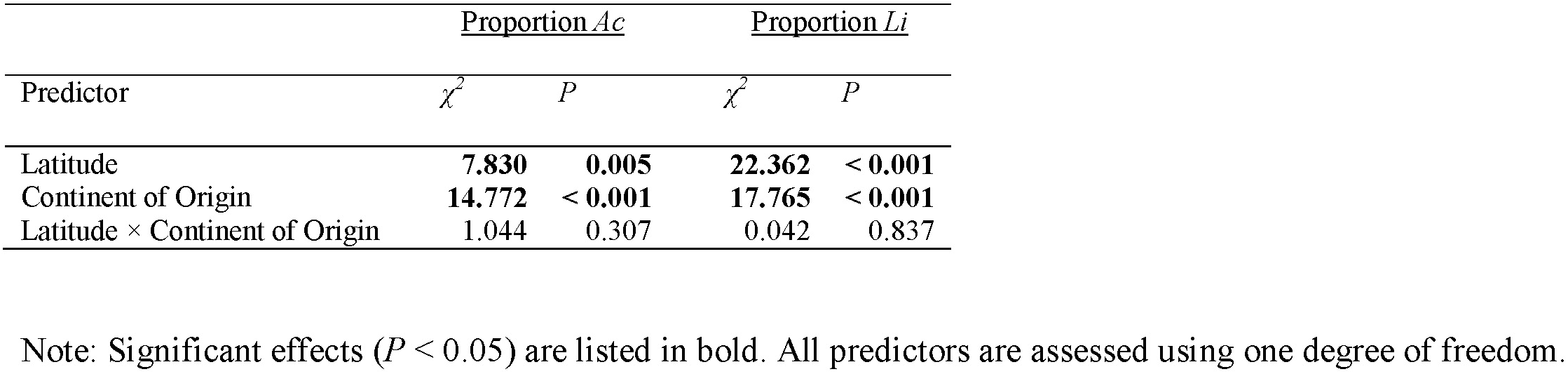
Results of mixed effects models assessing how latitude and continent of origin affected the proportion of *Ac* and *Li* alleles found in populations of *T. repens*.

| Predictor | <u>Proportion *Ac*</u> |  | <u>Proportion *Li*</u> |  |
| --- | --- | --- | --- | --- |
| | $\chi^2$ | *P* | $\chi^2$ | *P* |
| Latitude | **7.830** | **0.005** | **22.362** | **< 0.001** |
| Continent of Origin | **14.772** | **< 0.001** | **17.765** | **< 0.001** |
| Latitude × Continent of Origin | 1.044 | 0.307 | 0.042 | 0.837 |

Note: Significant effects ( $P < 0.05$ ) are listed in bold. All predictors are assessed using one degree of freedom.

**Table S11.** Results of mixed effects models assessing how cyanotype (AcLi, Acli, acLi, or acli), common garden location, and climatic distance affected the fitness of *T. repens*.

|  | <u>Plants Producing</u><br><u>Flowers</u> |  |  | <u>Plants Producing Seeds</u> |  |  | <u># of Flower Heads</u> |  |  | <u>Seed Set Mass</u> |  |  |
| --- | --- | --- | --- | --- | --- | --- | --- | --- | --- | --- | --- | --- |
| Predictor | $\chi^2$ | df | *P* | $\chi^2$ | df | *P* | $\chi^2$ | df | *P* | $\chi^2$ | df | *P* |
| Cyanotype (C) | **16.862** | **3** | **< 0.001** | **11.731** | **3** | **0.008** | 6.635 | 3 | 0.084 | 1.801 | 3 | 0.615 |
| Garden Location (L) | **29.547** | **3** | **< 0.001** | **43.214** | **3** | **< 0.001** | **125.477** | **3** | **< 0.001** | **19.881** | **3** | **< 0.001** |
| Climatic Distance (D) | **9.464** | **1** | **0.002** | **9.816** | **1** | **0.002** | **22.595** | **1** | **< 0.001** | **23.040** | **1** | **< 0.001** |
| C × L | 13.049 | 9 | 0.160 | 11.819 | 9 | 0.224 | 6.558 | 9 | 0.683 | 2.835 | 9 | 0.970 |
| C × D | 2.089 | 3 | 0.554 | 3.236 | 3 | 0.357 | 3.926 | 3 | 0.270 | 4.463 | 3 | 0.216 |
| C × L × D | 10.144 | 12 | 0.603 | 13.401 | 12 | 0.341 | 11.509 | 12 | 0.486 | 10.804 | 12 | 0.546 |

|  | <u>Survival</u> |  |  | <u>Maximum Plant Area</u> |  |  | <u>Growth Rate</u> |  |  | <u>Herbivory</u> |  |  |
| --- | --- | --- | --- | --- | --- | --- | --- | --- | --- | --- | --- | --- |
| Predictor | $\chi^2$ | df | *P* | $\chi^2$ | df | *P* | $\chi^2$ | df | *P* | $\chi^2$ | df | *P* |
| Cyanotype (C) | **19.889** | **3** | **< 0.001** | **9.841** | **3** | **0.020** | 0.381 | 3 | 0.944 | 4.956 | 3 | 0.175 |
| Garden Location (L) | **56.896** | **3** | **< 0.001** | **116.238** | **3** | **< 0.001** | **79.026** | **3** | **< 0.001** | **85.426** | **3** | **< 0.001** |
| Climatic Distance (D) | **3.903** | **1** | **0.048** | **6.887** | **1** | **0.009** | 3.835 | 1 | 0.050 | 0.237 | 1 | 0.626 |
| C × L | 8.506 | 9 | 0.484 | 7.049 | 9 | 0.632 | 6.369 | 9 | 0.703 | 3.326 | 9 | 0.950 |
| C × D | 1.723 | 3 | 0.632 | 2.726 | 3 | 0.436 | 1.251 | 3 | 0.741 | 3.724 | 3 | 0.293 |
| C × L × D | 6.019 | 12 | 0.915 | 6.575 | 12 | 0.884 | 19.085 | 12 | 0.087 | 8.448 | 12 | 0.749 |

Note: Significant effects ( $P < 0.05$ ) are listed in bold.

**Figure S1.**
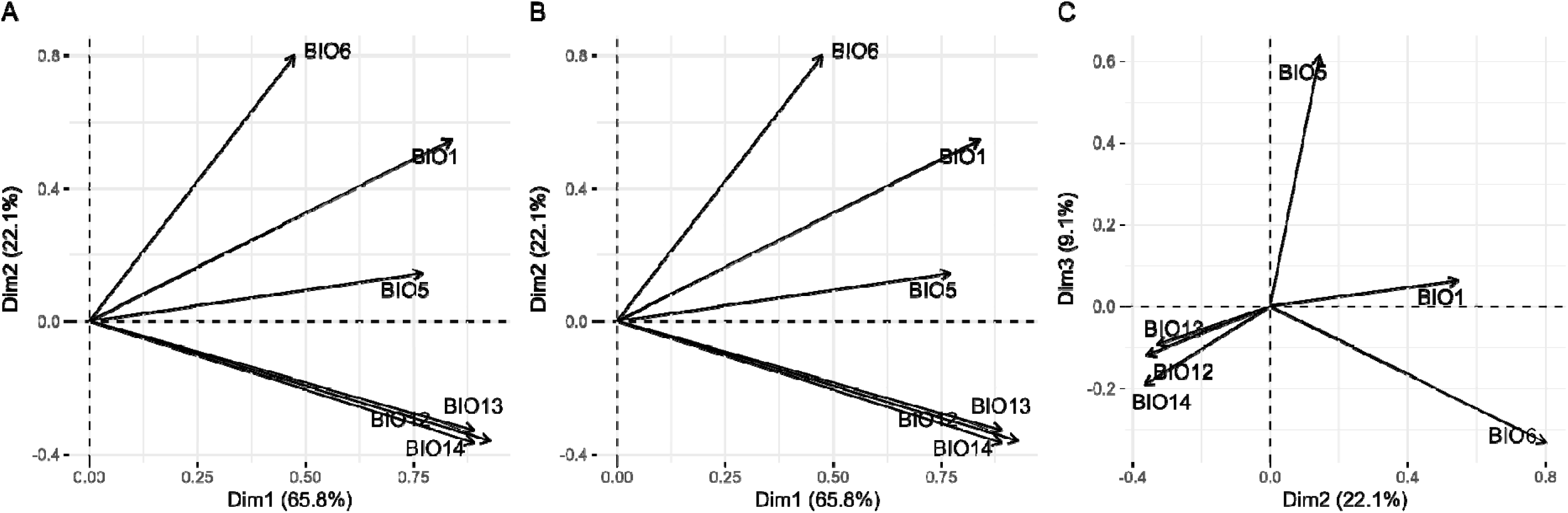
Eigenvectors of six bioclimatic variables (z-score standardized) on the three most influential dimensions of the principal component analysis used to determine climatic distance between each population of origin and each common garden. (**A**) Plot showing eigenvectors along dimensions 1 and 2. (**B**) Plot showing eigenvectors along dimension 1 and 3. (**C**) Plot showing eigenvectors along dimensions 2 and 3. Bioclimatic variables included in the principal component analysis are mean annual temperature (BIO1), maximum temperature of the warmest month (BIO5), minimum temperature of the coldest month (BIO6), annual precipitation (BIO12), precipitation of the wettest month (BIO13), and precipitation of the driest month (BIO14).

**Figure S2.**
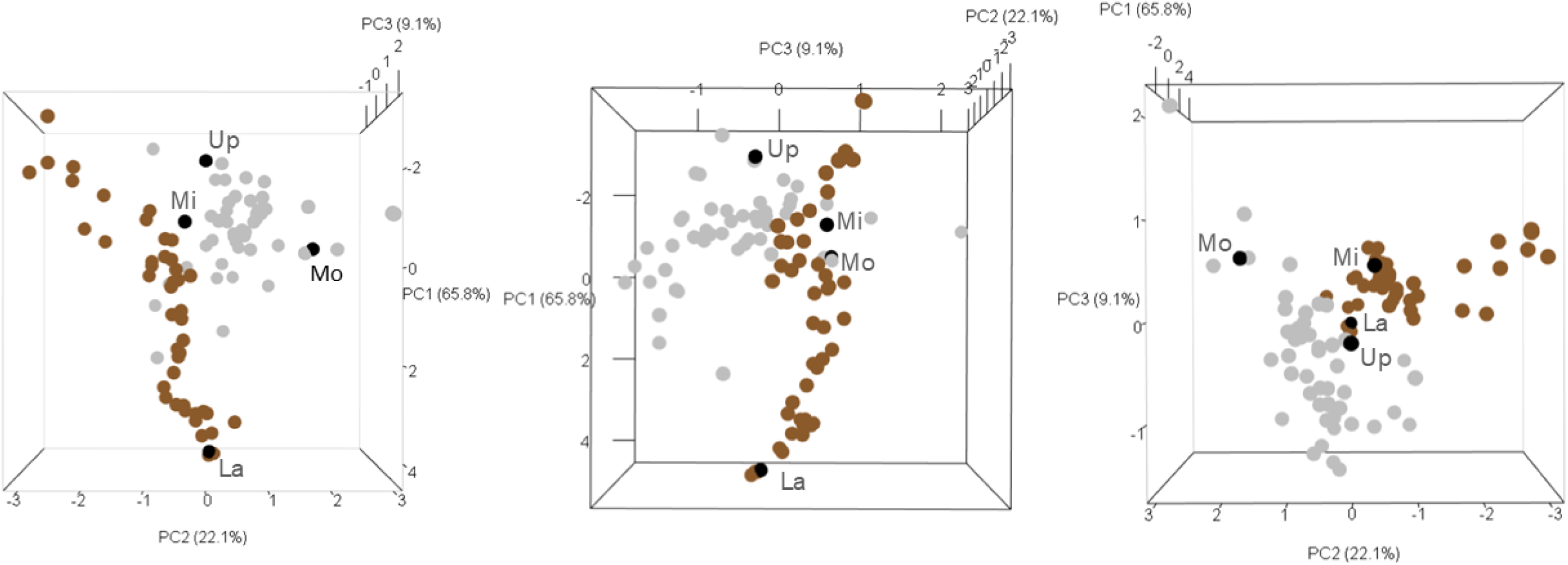
Results of a principal component analysis including six bioclimatic variables of interest, where each point represents the z-score standardized values on the three most influential dimensions (three-dimensional plot viewed from different angles) for each population from the native range (Europe, grey), each population from the introduced range (North America, brown) and each common garden location (black; Mississauga: Mi; Lafayette: La; Uppsala: Up; Montpellier: Mo). Weighted Euclidean distances were calculated between each population and each common garden using their PC1, PC2, and PC3 loading values as weights in order to determine a measure of climatic distance.

**Figure S3.**
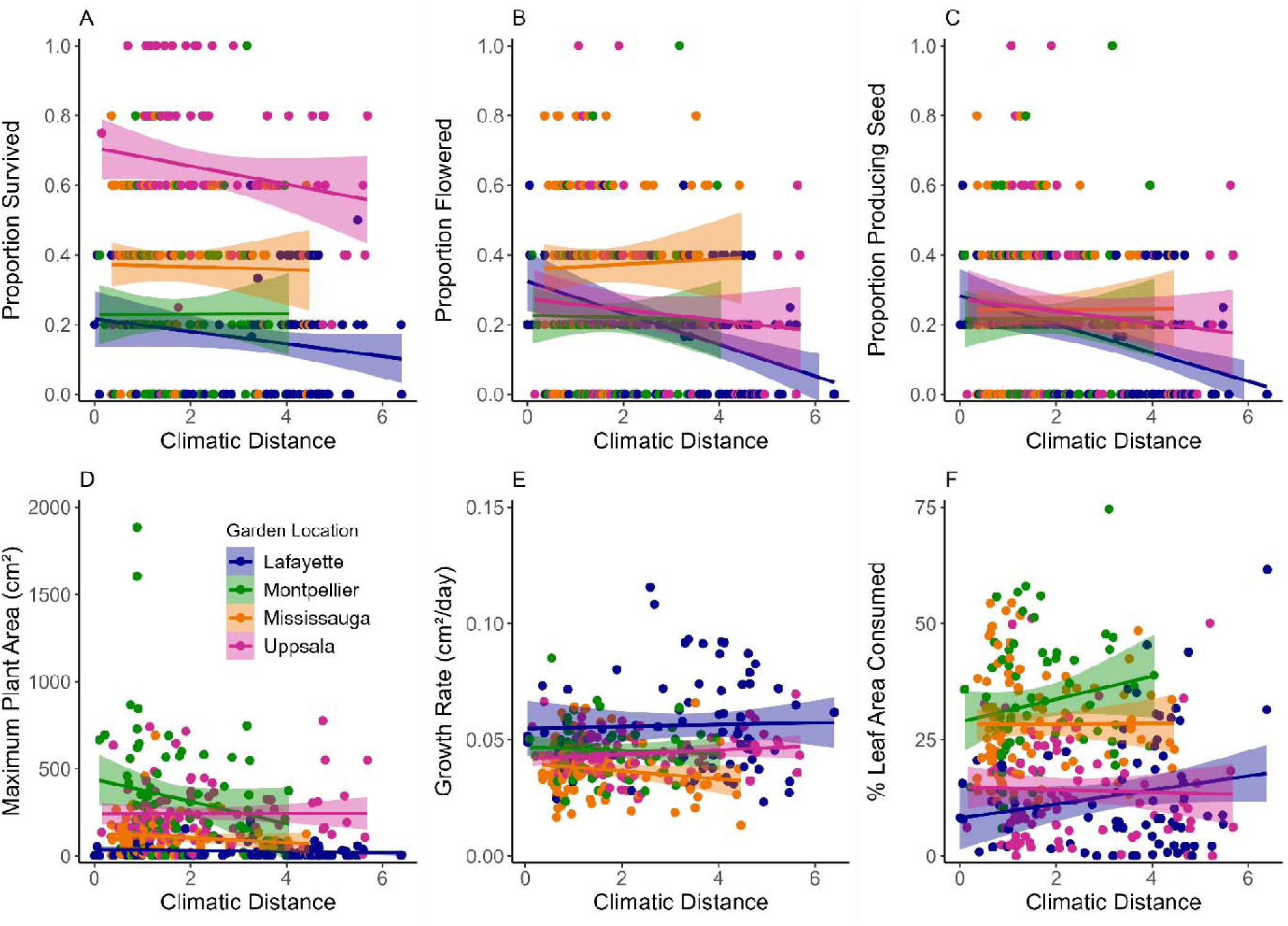
The effect of climatic distance and common garden location on fitness of *T. repens*. (**A**) Proportion of plants survived, (**B**) proportion of plants that produced flowers, (**C**) proportion of plants that produced seeds, (**D**) mean maximum plant area, (**E**) mean growth rate and (**F**) mean herbivory (measured as the percentage of leaf area consumed) for *T. repens* populations planted in four common gardens (Lafayette in navy, Montpellier in green, Mississauga in orange, and Uppsala in magenta). Shaded areas represent 95% confidence intervals around each line.

**Figure S4.**
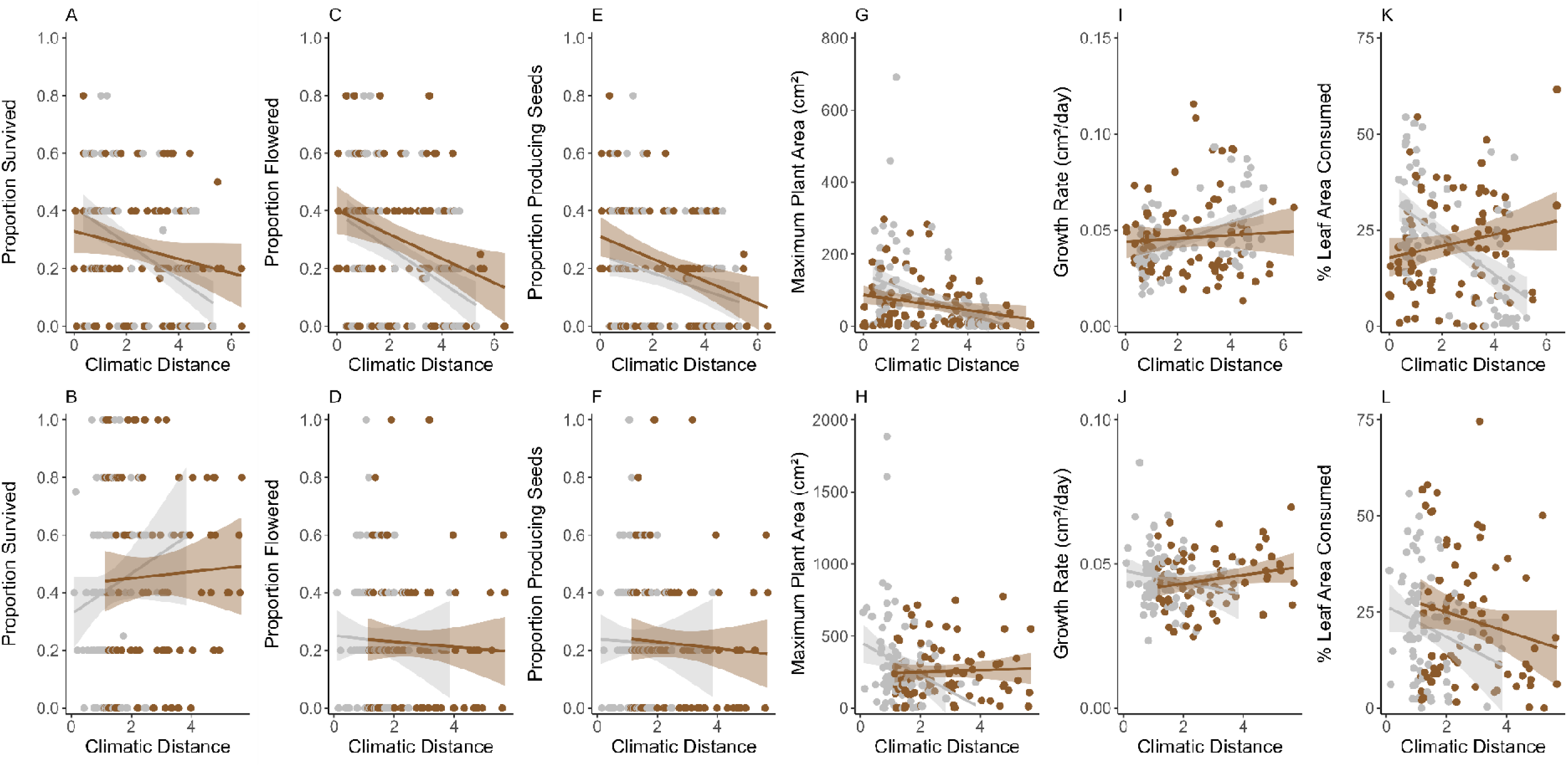
The effect of climatic distance on the proportion of *T. repens* survived when originating from the introduced (North America, orange) and native ranges (Europe, pink) and transplanted into common gardens in the (**A**) introduced and (**B**) native ranges. The effect of climatic distance on the proportion of plants flowered when originating from the introduced and native ranges and transplanted into the (**C**) introduced and (**D**) native ranges. The effect of climatic distance on the proportion of plants producing seeds when originating from the introduced and native ranges and transplanted into the (**E**) introduced and (**F**) native ranges. The effect of climatic distance on the maximum area of plants originating from the introduced and native ranges and transplanted into the (**G**) introduced and (**H**) native ranges. The effect of climatic distance on growth rate of plants originating from the introduced and native ranges and transplanted into the (**I**) introduced and (**J**) native ranges. The effect of climatic distance on herbivory of plants originating from the introduced and native ranges and transplanted into the (**K**) introduced and (**L**) native ranges. Points represent mean values for each population within each garden. Shaded areas represent 95% confidence intervals around each line.

**Figure S5.**
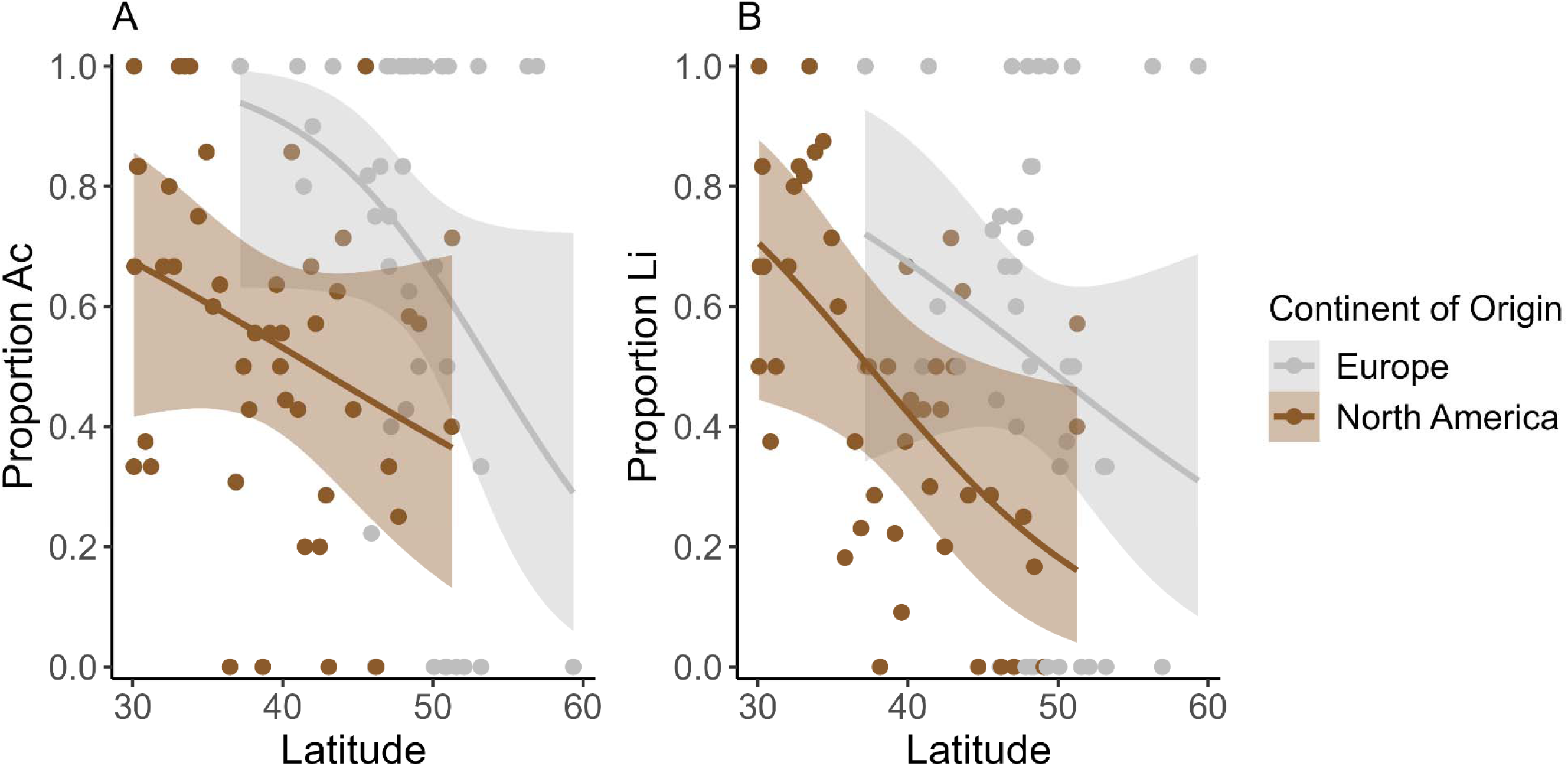
(**A**) The proportion of *T. repens* plants possessing at least one dominant *Ac* allele per sampled population in either the native range (Europe, pink) or introduced range (North America, orange) as a function of the latitude of origin of each population. (**B**) The proportion of *T. repens* plants possessing at least one dominant *Li* allele per sampled population in either the native or introduced range as a function of the latitude of origin of each population. Shaded areas represent 95% confidence intervals around each line.

**Figure S6.**
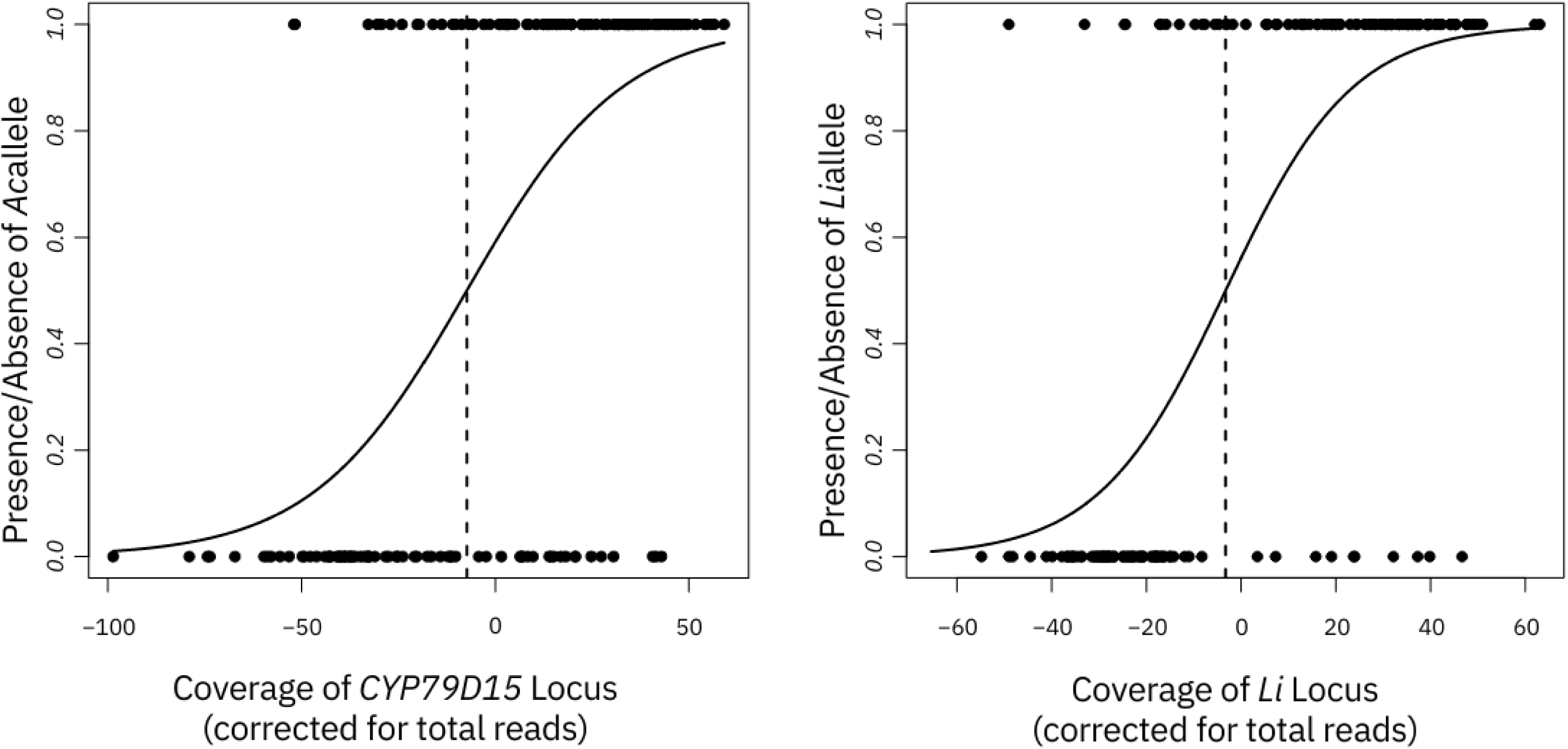
Associations between inferred genomic coverage of cyanogenesis genes and Feigl-Anger cyanogenesis phenotypes. Coverage of cyanogenesis genes is inferred from low coverage whole genome sequencing reads found within loci associated with the *Ac*/*ac* and *Li*/*li* genes. The x-axis is corrected for the total number of reads because samples with few reads may not have reads within these regions by chance (i.e., not because the plant possessed the associated gene deletion). Lines represents generalized linear models fit with a binomial distribution.

## Supplementary Methods

### DNA Extraction and Library Preparation

Our DNA extraction and library construction have been previously described (Battley et al., 2024). Briefly, we collected 1-3 trifoliate leaves per plant and either stored tissue at −80°C until extraction at University of Louisiana, Lafayette (ULL) or lyophilized tissue prior to shipping to ULL. DNA was extracted with a modified cetyltrimethylammonium bromide (CTAB) procedure using increased concentrations of CTAB and 2-β-mercaptoethanol (Doyle and Doyle, 1987; Saeidi et al., 2018). DNA concentrations were quantified and diluted to 3 ng/uL before undergoing a modified Nextera dual-index library preparation (Therkildsen and Palumbi, 2017). We used Nextera XT Index Kit (Illumina; San Diego, CA, USA) for barcoding libraries. Each library was quantified and concentrations normalized before pooling of 48 samples with unique barcodes into a single tube. Libraries were sequenced on a HiSeq X Ten platform by Novogene (Sacramento, CA, USA) using paired end 150 bp reads. In total, 656 samples (Lafayette, Louisiana, USA = 138; Mississauga, Ontario, Canada = 327; Montpellier, France = 111; Uppsala, Sweden = 79) were sequenced across 14 sequencing lanes.

### Bioinformatics Pipeline

For each sample, adapter sequences and poly-g tails were trimmed from reads using *fastp* (Chen et al., 2018). An FM-index was constructed for the *Trifolium repens* reference genome (Santangelo et al., 2023) using the bwa indexing algorithm with default settings (Li and Durbin, 2009). Reads were then aligned to the reference genome using bwa mem (Li and Durbin, 2009). SAM files were converted to BAM file format, coordinate sorted, and indexed using SAMtools (Li et al., 2009). To find the genomic coordinates of the *Ac*/ac and *Li/li* loci in the *T. repens* reference genome, we took 5 sample sequences of the *CYP79D15* locus (the *Ac/ac* gene; Olsen et al. 2008) and *Li* locus (*Li/li* gene; Olsen et al., 2007) from the NCBI database and compared nucleotide similarity of NCBI samples with our reference genome using the basic local alignment search tool (BLAST; Camacho et al., 2009). We found 99% similarity between 5 NCBI nucleotide sequences of the *Li* gene with Chr04_Pall:28511241 - 28515079. For *CYP79D15*, we found 99% sequence similarity at sites Chr02_Occ:9222822 – 9224484 and Chr02_Occ: 9536228 – 9537890. Constraining our search to these regions, we calculated the number of reads aligning to these regions using the ‘view -c’ command and calculated read coverage using the ‘coverage’ command within SAMtools (Li et al., 2009). This resulted in number of reads and read coverage data for 586 samples (Lafayette, Louisiana, USA = 99; Mississauga, Ontario, Canada = 302; Montpellier, France = 107; Uppsala, Sweden = 78) after removing samples that sequenced poorly (< 100 Mb of read data).

### Statistics

Our low coverage whole genome sequencing approach for direct determination of genotypes at the *Ac/ac* or *Li/li* genes (referred to below as cyanotypes). We took a statistical genomic approach to determining genotypes and evaluating potential type I and type II error in our cyanotypes. Since variation at *Ac/ac* and *Li/li* is caused by deletions, we used read coverage within the underlying genomic regions as evidence for the presence of the *Ac* or *Li* alleles. We first removed any samples with < 500,000 reads (20 samples). We use read coverage of each locus rather than number of reads as number of reads is likely more influenced by variable gene deletion sizes and misalignment of homeologous sequences. Indeed, our downstream analyses suggest the number of reads metric produces higher error than coverage in genotype estimates. For each locus, we first correct our coverage values for variation in sequencing effort using a general linear model with the *lm* function in R Version 4.2.2 (2022) to associate coverage with total number of reads across the entire genome. The residuals from this model reflect effort-corrected coverage of a locus. We then use a set of individuals (N = 178) whose cyanotypes were previously determined by Feigl-Anger assays to model the relationship between coverage at each locus and the presence of at least one dominant allele (*Ac* or *Li*; Feigl and Anger, 1966). Separate univariate generalized linear models implemented using the *glm* function with a binomial family and logit link were conducted for each locus with presence or absence of the dominant allele assessed via Feigl-Anger assay as the response variable and the residual of coverage model as the predictor variable. Statistical significance of these associations was assessed via a Wald statistic implemented via the *Anova* function in the *car* package (version 3.1-2; Fox et al., 2013).

These models were highly significant for each locus (*Ac*: *χ*^2^ = 77.3, df = 1, *P* < 2.2e-16; *Li*: *χ*^2^ = 110.6, df = 1, *P* < 2.2e-16; **Figure S6**), indicating that coverage was a valuable predictor of cyanotype. From our predicted models, we calculated the inflection point of the model for each locus and used this value to predict the presence or absence of *Ac* and *Li* alleles for every sample. Plants with higher values than the inflection point that the model estimate for each locus were called as having either the *Ac* or *Li* allele present. Using this method to estimate cyanotypes of samples with known cyanotypes via FA-assay, we find 142/178 genotypes were correctly determined for the *Ac/ac* gene (79.8%) and 154/178 for the *Li/li* gene (86.5%). Errors were skewed toward type II error (false negatives) as would be expected with our low coverage whole genome sequencing approach. We also note that Feigl-Anger assays have some inherent error, especially when conducted on small amount of tissue from field-grown plants. We are unable to attribute the differences between approaches to error in either of the assays and thus just note the substantial concordance.

